# Agonist efficacy at the β_2_AR is driven by agonist-induced differences in receptor affinity for the G_s_ protein, not ligand binding kinetics

**DOI:** 10.1101/2024.01.05.574357

**Authors:** Clare R. Harwood, David A. Sykes, Theo Redfern-Nichols, Graham Ladds, Stephen J. Briddon, Dmitry B. Veprintsev

**Author notes:** **Correspondence:** Stephen J. Briddon and Dmitry B. Veprintsev.

## Abstract

**Introduction:** The β_2_-adrenoceptor (β_2_AR) is a class A G protein-coupled receptor (GPCR). It is therapeutically relevant in asthma, whereby β_2_AR agonists relieve bronchoconstriction. The β_2_AR is a prototypical GPCR for structural and biophysical studies. However, the molecular basis of agonist efficacy at the β_2_AR is not understood. We hypothesized that the kinetics of ligand binding and GPCR-G protein interactions could play a role in ligand efficacy. We characterised the molecular pharmacology of a range of β_2_AR agonists and examined the correlation between ligand and mini-G_s_ binding kinetics and efficacy.

**Methods:** We used a Time-resolved Fluorescence Resonance Energy Transfer (TR-FRET) based competition ligand binding assay to measure the affinity and residence times of a range of β_2_AR agonists binding to the human β_2_AR. TR-FRET between Lumi4-Tb^3+^ on the N terminus of the β_2_AR and fluorescent CA200693 (S)-propranolol-green was measured using a PHERAstar FSX. The ability of these β_2_AR agonists to activate the heterotrimeric G_s_ protein was measured using the CASE G_s_ protein biosensor. This assay senses a reduction in NanoBRET between the nano-luciferase (nLuc) donor on the Gα subunit and Venus acceptor on the Gγ, on receptor activation, quantified using the operational model of agonism. NanoBRET was also used to measure interactions between DDM solubilised β_2_AR-nLuc and purified Venus-mini-G_s_. A large excess of unlabelled mini-G_s_ was used to dissociate the β_2_AR-nLuc: Venus-mini-G_s_ complex.

**Results:** Characterisation of the molecular pharmacology of seven β_2_AR agonists showed a broad range of ligand binding affinities (p*K*_i_ = 4.4 ± 0.09 to 9.2 ± 0.08) and kinetics parameters. There was no correlation between ligand residence times and their ability (log ***τ***) to activate the G_s_ protein (R^2^=0.26, p=0.29). However, there were statistically significant differences in the association rate (*k*_on (fast)_) (3.36±0.64×10^5^ to 9.19± 0.42×10^5^) and affinity (*K*_d_) values of mini-G_s_ binding to the agonist-β_2_AR complex (p*K*_d_ =6.0 to 6.7). Both an increase in ligand driven mini-G_s_ *k*_on(fast)_ rate and associated increase in mini-G_s_ p*K*_d_ for the receptor, were moderately correlated with efficacy (log ***τ***) (R^2^ =0.58 and R^2^ =0.50 respectively).

**Conclusions:** These data support a model in which agonists of increased efficacy cause the β_2_AR to adopt a conformation that is more likely to recruit G protein. Conversely, these data did not support a role for agonist binding kinetics in the molecular basis of efficacy.

## Introduction

G protein-coupled receptors (GPCRs) are the largest family of membrane proteins within the human genome and responsible for modulating a broad range of hormonal, neurological and immune responses. GPCR-directed therapeutics currently target over 100 diverse receptors and represent 34% of all US food and drug administration (FDA) approved drugs, making them the most widely targeted receptors(Hauser et al., 2017). Despite their therapeutic importance, the molecular basis of ligand efficacy, the ability of a drug to produce an effect at GPCRs, is not fully understood. It is hoped that an increased understanding of the molecular basis of efficacy would aid more rational drug design.

Several studies have implicated the role of ligand residence time in the molecular basis of efficacy at GPCRs. For example, a positive correlation has been shown between the efficacy of seven agonists at the muscarinic M3 receptor, and ten agonists at the adenosine A_2A_ receptor and their ligand residence time (Sykes et al., 2009; Guo et al., 2012). Conversely, no correlation between efficacy and residency time was shown for ligands at the adenosine A_1_ receptor (Louvel et al., 2014).

Biophysical studies have shown that agonists shift the receptor conformational landscape in favor of a unique active conformation, compared to the unliganded state (Deupi and Kobilka, 2010; Mary et al., 2012; Nygaard et al., 2013), however, how conformational differences in a population translates to greater or lesser signaling responses remains to be fully elucidated. Structural studies have found little differences in GPCR conformations adopted by ligand-bound GPCR-G-proteins complexes (Masureel et al., 2018; Zhang et al., 2020). However, using Nuclear Magnetic Resonance (NMR), Lui and colleagues (Liu et al., 2012) showed efficacy-dependent differences in the conformational state of β_2_AR bound to different agonists, prior to G protein binding. Similar results have been seen for the β_1_AR (Grahl et al., 2020) and A_2A_ (Ye et al., 2016). Alternatively, some studies (Nikolaev et al., 2006; Gregorio et al., 2017) show correlations between ligand efficacy and the rate of GPCR and G protein activation, suggesting a role for G protein binding kinetics in the molecular basis of efficacy.

Consequentially, we aimed to delineate the roles of ligand binding and receptor-G protein binding kinetics in agonist efficacy. We have focused on the β_2_-adrenoceptor (β_2_AR), a prototypical class A GPCR, which is one of the structurally, functionally, and therapeutically best characterised GPCRs. The β_2_AR is also an essential target in the treatment of asthma and COPD, and as consequence a wide range of clinically used agonists of varying efficacies have been produced to target the β_2_AR, which could be utilised in this study.

To this end, we characterised the ligand binding affinities, residence times and efficacy at the level of heterotrimeric G_s_ protein activation of seven agonists at the β_2_AR. Additionally, we investigated the binding kinetics and affinity of fluorescently labelled mini-G_s_ proteins for the β_2_AR in complex with these seven agonists. As previously suggested, we found no correlation been agonist binding kinetics and efficacy (Sykes and Charlton, 2012). Alternatively, we found a trend between ligand-induced differences in G protein binding kinetics and affinity for the agonist bound β_2_AR and an agonists ability (efficacy) to activate the heterotrimeric G_s_ protein. These findings suggest differences in initial agonist-GPCR conformations could be key to understanding the molecular basis of efficacy of these seven β_2_AR agonists.

## Materials and Methods

### Materials

The T-REx™-293 cell line was obtained from Invitrogen (CA, U.S.A). T75 and T175 mammalian cell culture flasks were purchased from Fisher scientific (Loughborough, UK). All cell culture reagents, including Phosphate Buffered Saline (PBS) and Fetal Calf Serum (FCS) were purchased from Sigma Aldrich (Gillingham, UK), except for blastocidin which was obtained from Gibco™ (MA, U.S.A) and zeocin™. Tag-lite SNAP-Lumi4-Tb labelling reagent and LABMED buffer was purchased from Cisbio (Codolet, France). Polyethylenimine (PEI) (25kDa) was obtained from Polysciences Inc (PA, U.S.A), and CellStar 96 well tissue culture plates from Greiner Bio-One (Kremsmünster, Austria). *n*-Dodecyl β-D-maltoside (DDM) were obtained from Anatrace (OH, U.S.A). HisTrap FF crude 5mL columns were obtained from GE Healthcare (IL, U.S.A). Vivaspins protein concentrators were obtained from Sartorious (Göttingen, Germany). Slide-a-Lyzer dialysis cassettes, NuPAGE LDS sample buffer, NuPAGE 4-12% Bis-Tris 15 x 1.0mm well gels, NuPAGE MOPs SDS running buffer, Pageruler pre-stained protein ladder and BODIPY F-L-cystine dye were all obtained from Thermofisher (MA, U.S.A). β_2_AR antagonist [(S)-propranolol-green] (CA200693), was obtained from CellAura, UK, and supplied by Hello Bio, (Bristol, U.K) (s)-(-)- Propranolol hydrochloride, salmeterol were obtained from Tocris, (Bristol, U.K). Formoterol hemifurmate from APExBIO (TX, U.S.A), and BI-167-107 from Boehringer Ingelheim (Ingelheim, Germany). (±)-epinephrine hydrochloride, noradrenaline, salbutamol hemisulfate and isoprenaline hydrochloride were purchased from Sigma Aldrich (Gillingham, UK). Nano-Glo luciferase substrate was obtained from Promega (WI, U.S.A). All other chemicals were purchased from Sigma Aldrich (Gillingham, UK).

### Instruments and software

BMG PHERAstar FSX plate reader (BMG Labtech, Offenburg, Germany), fitted with BRET1 plus, TRF 337 520/620, TRF and LUM 550LP/450-80 optic modules and MARS software were purchased from BMG Labtech (Offenburg, Germany). GraphPad Prism 9 was purchased from GraphPad Software, (San Diego, U.S.A.). Microsoft Excel^TM^ XP was purchased from Microsoft (Washington, U.S.).

### Methods Molecular biology

The construct pcDNA4TO-TwinStrep (TS)-SNAP-β_2_AR was generated by amplification of the SNAP and β_2_AR sequences of the pSNAPf-ADRB2 plasmid (NEB) and inserted into pcDNA4TO-TS using Gibson assembly (Heydenreich et al., 2017). pcDNA4TO-TS-SNAP-β_2_AR-nLuc was generated by Dr Brad Hoare by amplification of pcDNA4TO-TS-SNAP-β_2_AR and nanoLuc, with insertion of nanoLuc into pcDNA4TO-TS-SNAP-β_2_AR via Gibson assembly. Both constructs used a signal peptide based on the 5HT_3A_ receptor to increase protein folding and expression. The CASE G_s_ protein construct is that designed and optimised by the Schulte lab (Schihada et al., 2021) and were obtained from Addgene. Mammalian mini-G_s_ constructs were a kind gift from Nevin Lambert (Wan et al., 2018). For bacterial expression of Venus-mini-G_s_ and mini-G_s,_ protein encoding DNA sequences were amplified from corresponding mammalian constructs and inserted into the pJ411 vector containing MKK-HIS10-TEV N-terminal tag (Sun et al., 2015), via Gibson assembly to give the constructs MKK-HIS10-TEV-mini-Gs, and MKK-HIS10-TEV-Venus-mini-G_s_.

### Transfection and mammalian cell culture

pcDNA4TO-TS-SNAP-β_2_AR or pcDNA4TO-TS-SNAP-β_2_AR-nLuc were stably transfected into T-Rex^TM^-293 cells (Invitrogen) using polyethylenimine (PEI). A stable mixed population was selected by resistance to 5 µg/mL blasticidin and 20 µg/mL zeocin. Stable cell lines were maintained in high glucose DMEM (Sigma D6429) with 10% foetal bovine serum (FBS), 5μg/μL blasticidin and 20μg/μL zeocin at 37°C and a humidified atmosphere of 5% CO_2._ When ∼70% confluent TS-SNAP-β_2_AR or TS-SNAP-β_2_AR-nLuc expression was induced with 1μg/mL tetracycline. Cells were left to express for 50hrs before harvesting for assays. The T-Rex^TM^-293 pcDNA4TO-TS-SNAP-β_2_AR-CASE G_s_ stable cell line was generated by stably transfecting the CASE G_s_ constructs into the T-Rex^TM^-293 pcDNA4TO-TS-SNAP-β_2_AR using PEI. A mixed population stable cell line was generated by selection with 500 µg/mL G418 and then a single colony population generated via FACS.

### CASE-Gs activation assays

For CASE-G_s_ activation assays, a single population of T-Rex^TM^-293 stably expressing pcDNA4TO-TS-SNAP-β_2_AR and CASE G_s_ was plated at 50,000 cells/well in 96 well plates in a volume of 100μL and induced for 48hrs with 1μg/mL tetracycline at 37°C and 5% CO_2_. Plates were washed twice with 100μL/well HBSS prior to addition of 80μL/well assay buffer (HBSS (Sigma H8264) + 0.1% BSA). 10μL of x10 furimazine diluted in assay buffer was added to each well, to give a final concentration of 8μM and plates were incubated at 37°C and 5% CO_2_ for 20mins. A white back seal was placed on the underside of the plate and luminescence was read on a PHERAstar FSX using 450-80/550LP module for 3 mins to establish a baseline BRET signal. The plate reader was then paused and 10μL of x10 ligand dilutions were added accordingly.

### Labelling TS-SNAP-β2AR with Terbium cryptate for TR-FRET ligand binding assays

Media was aspirated from T175 flasks and adherent cells washed twice at room temperature with Phosphate Buffered Saline (PBS). Adherent cells were labelled with 100nM SNAP-Lumi4-Tb labelling reagent in Labmed buffer (Cisbio, UK) for 1 hr at 37°C and 5% CO_2_. Post incubation cells were washed twice more with PBS and detached with 5mL non-enzymatic cell dissociation solution (Sigma, UK). Cells were pelleted by centrifugation for 10 min at 1000*xg*, and cell pellets frozen at -80°C.

### Solubilisation of the TS-SNAP-β2AR or TS-SNAP-β2AR-nLuc

TS-SNAP-β_2_AR or TS-SNAP-β_2_AR-nLuc were solubilised from stably transfected T-Rex^TM^-293 cell membranes, generated as described previously (Harwood et al., 2021). Solubilisation took place using 1% DDM (w/v) in 20mM HEPES, 5% (v/v) glycerol, and 150mM NaCl, pH 8 at 4°C for 2-3 hrs. Samples were clarified by ultracentrifugation at 4°C for 1hr at 100,000*xg*.

### TR-FRET competition ligand binding

TR-FRET between the donor Lumi4-Tb and the fluorescent acceptor CA200693 (S)-propranolol-green (75nM) was measured by exciting at 337nm and quantifying emission at 520nm and 620nm using a PHERAStar FSX (BMG Labtech) and TRF 337 520/620 module (BMG Labtech). Assay buffer consisted of 20mM HEPES, 5% glycerol, 150mM NaCl, 0.1% DDM and 0.5% BSA, pH 7.4. All binding assays used a final concentration of 1% Dimethyl sulfoxide (DMSO), assay volume of 30µL, 384 well OptiPlates (PerkinElmer, US) and 1µM alprenolol to determine non-specific binding (NSB).

Receptors were added last, and plates were incubated at room temperature for 20-40 mins prior to reading, depending on the kinetics of the ligand.

### Production of mini-Gs

His-TEV-Venus-mini-G_s_ and His-TEV-mini-G_s_ were expressed in NiCo21(DE3) *E.*coli, cultured in Terrific Broth (Gibco). 1L cultures were induced with 1mM Isopropyl β-D-1-thiogalactopyranoside (IPTG) at OD=0.6 and incubated for a further 20 hrs at 20°C and 225 RPM. Pellets from 1L cultures were thawed on ice, and resuspended in 50mL lysis buffer (20mM HEPES, pH 7.5, 500 mM NaCl, 40 mM imidazole, 10% glycerol, 8 mM β-mercaptoethanol (BME), 1 μM guanosine diphosphate (GDP), cOmplete protease inhibitors (Roche), DNAase I, and lysozyme using a dounce homogeniser. Lysis took place on ice via sonication, using a Vibra cell probe sonicator with 5 x 10 second pulses, 30 seconds apart. Lysate was loaded onto HisTrap FF crude 5mL column, using using ÄKTA^TM^ start protein purification system at a flow rate of 5 mL/min. System and column had been equilibrated with 10 column volumes (CV) buffer A (20mM HEPES, 500mM NaCl, 40mM imidazole, 10% glycerol, 8mM BME, 1μM GDP. Unbound protein was washed out with 10 CV buffer A. Bound protein was then eluted over an 8 CV gradient of 0-100% buffer A to B at a flow rate of 5 mL/min. (Buffer B= 20mM HEPES, 500mM NaCl, 400mM imidazole, 10% glycerol, 8mM BME and 1μM GDP). The prescence of His-TEV-Venus-mini-G_s_ and His-TEV-mini-G_s_ was confirmed by SDS-Page analysis and InstantBlue stain for protein. Pooled elution fractions were then concentrated using 10,000 or 30,000 molecular weight cut-off (MWCO) Vivaspin protein concentrators by centrifugation at 3000*xg* and 4° C for 15 min intervals over 2-3 hours. Protein was exchanged into assay buffer using slide-A-lyzer^TM^ 10,000 or 30,000 MWCO dialysis cassettes for untagged and Venus-tagged mini-Gs protein samples, respectively. Dialysis took place overnight at 4°C under constant stirring. Assay buffer consisted of 20mM HEPES, 150mM NaCl, 10% glycerol, 8mM BME and 1μM GDP. Purified mini-Gs protein was flash frozen using liquid nitrogen and stored at -80°C.

### In-solution TS-SNAP-β2AR-Venus-mini-Gs nanoBRET binding assays

20mM HEPES, 150mM NaCl, 10% glycerol, 1μM GDP, 8mM BME, 0.5% BSA and 0.1% ascorbic acid pH 7.4 was used as the assay buffer in all in-solution nanoBRET assays. For recruitment assays, varying concentrations of β_2_AR agonists were used to recruit Venus-mini-G_s_ to the DDM-TS-SNAP-β_2_AR, from an excess (1μM) of Venus-mini-G_s_. Assays were run in 20μL volumes in white 384 well proxiplates. 25μM unlabelled mini-G_s_ was used to define specific binding of the Venus-mini-G_s_ to the TS-SNAP-β_2_AR-nLuc receptors. Receptor, ligand and mini-G_s_ proteins were added to plate and incubated for 80 mins at room temperature, 8μM furimazine final concentration was added to plate and incubated for a further 10 mins before reading on PHERAstar FSX using 450-80/550LP module. For kinetic assays in which the affinity of Venus-mini-G_s_ for the agonist bound TS-SNAP-β2AR-nLuc receptors was measured over time, assays were run in 20μL volumes in white 384 well proxiplates. Varying concentrations (1.4 to 3000 nM) of Venus-mini-G_s_ were added to plates with either buffer or 30μM mini-G_s_ to define total and non-specific binding, respectively. DDM solubilised receptors were incubated with saturating concentrations of selected β_2_AR agonists for 40 mins, and 4X (32μM) furimazine for 10 mins, prior to addition to plate. Receptor was added to the plate offline, mixed up and down rapidly with a matrix pipette and read immediately on PHERAstar FSX as described above. After reading for 20 mins, the reader was paused and 2μL of 333μM mini-G_s_ was added to total wells to dissociate, plate was read for a further 20 mins (2μL buffer was added to NSB wells).

### Mathematical modelling

Here we have used our previously described ordinary differential model (ODE) of the cubic ternary complex model (Weiss et al., 1996) containing additional reactions to simulate the G protein activation cycle (Woodroffe et al., 2009; Bridge et al., 2018). The model, encoded in COPASI (Hoops et al., 2006), includes ligand binding, receptor activation, G protein binding and the G protein cycle, whereby the model output is activated G protein Gα_GTP_ and receptor occupancy (Bridge et al., 2018). Prior to addition of ligand, we first compute the system for 10^6^ seconds. To enable simulation of the data the co-operativity factor β (see SF6; table ST4) was varied and simulations computed. Steady state was reached after 5 minutes and outputs are shown after 10 minutes.

### Data analysis

All non-linear regression and statistical analysis were performed in GraphPad Prism 9. Where multiple replicates were combined, such as TR-FRET equilibrium binding curves and mini-G_s_ equilibrium recruitment curves shown below, data points for each replicate was normalised to the maximum value obtained for each ligand in each experiment. Competition ligand binding data were fitted to a one site model (equation 1).

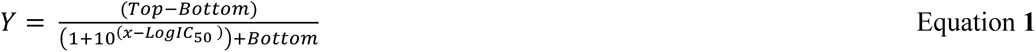

Where: Y = binding of tracer, x = Log[ligand], IC_50_ = the concentration of competing ligand which displaces 50% of radioligand specific binding.

*K*_i_ values were obtained using the Cheng-Prusoff equation (equation 2). Final *K*_i_ values were taken as a mean of individual experiments.

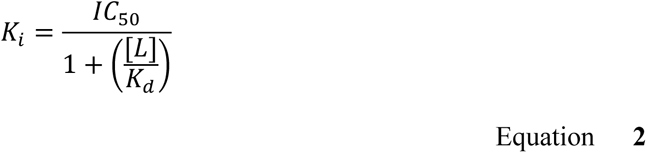

Where: Ki = the inhibition constant of the unlabelled ligand, [L] = concentration of labelled ligand, Kd = the equilibrium dissociation constant of the labelled ligand.

CASE G_s_ activation data from individual replicates were fitted to the operational model with *K*_A_ values fixed to those obtained in ligand binding studies (*K*_i_).

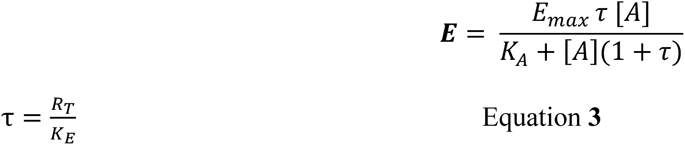

Where: τ− = the transducer ratio, K_A_ = the equilibrium association constant of the agonist, [A] = concentration of agonist, R_T_ = total receptor concentration, K_E_ = the concentration of agonist-receptor complex required for half maximal response.

Saturation binding curves for Venus-mini-G_s_ binding the agonist-TS-SNAP-β_2_AR-nLuc were fitted to a one site specific binding model according to equation 4. Final *K*_d_ values were taken as an average of *K*_d_ values from individual specific curve fits.

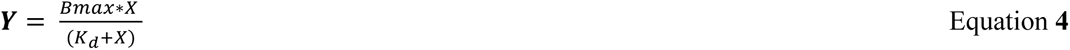

Where: Y = specific binding, *K*_d_ = the equilibrium dissociation constant of the labelled ligand, x= [Venus-mini-G_s_].

For kinetic studies on Venus-mini-G_s_ binding to TS-SNAP-β_2_AR-nLuc, Venus-mini-G_s_ association to the receptor was fitted to a two-site exponential association model and dissociation to a one phase exponential decay model, according to equations **5** and **7**.

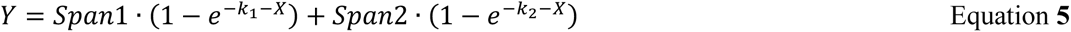

Where, _X=_ time, Span = difference between top and bottom for phases 1 and 2 respectively.

*k*_obs_ plots for *k*_fast_ values obtained from equation 5 were fitted to a simple linear regression model according to equation 6 to obtain the *k*_on_ of *k*_fast._

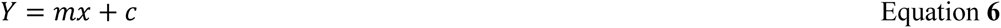

Where: m = slope or *k*_on_, c = intercept of y or *k*_off,_ *x*= concentration of Venus-mini-G_s_

For the above analysis the intercept was fixed to *k*_off_ values measured experimentally and obtained via equation 7.

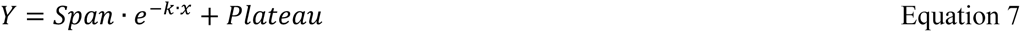

Specific binding data for the dissociation of Venus-mini-G_s_ from the agonist bound DDM-β_2_AR complex was fitted to a one phase exponential decay model, according to equation 7. Where Y = specific binding, span = difference between top and bottom, *k*= the rate constant, and plateau = y value at infinite times.

### Statistical analysis

Comparison of p*K*_d_, p*K*_i,_ *k_off_*, *k*_on,_ and *1″* values were made using a one-way Analysis Of Variance (ANOVA) test and Tukey’s post hoc multiple comparison test. Statistical comparison of p*EC_50_* values obtained from mini-G_s_ recruitment assays and *K*_i_ ligand binding values for each ligand were made using unpaired t-tests. A Pearson’s correlation coefficient was used to investigate correlations between CASE-G_s_ activation, *τ* values and relative time to reach equilibrium (IC_50 1min_/IC_50 Equilibrium_) and between CASE-G_s_ activation 1″ values and mini-G_s_ binding *k*_on_ and *k*_off_ values. All statistical analysis was completed in GraphPad Prism 9 and p<0.05 was considered statistically significant.

## Results

### Characterisation of the molecular pharmacology of seven β2AR agonists

We chose seven β_2_AR agonists anticipated to have a diverse range of efficacies, affinities and ligand binding kinetics. We characterised these parameters using a TR-FRET ligand binding assay and BRET-based G_s_ protein activation assay. **Figure 1A** shows competition binding curves for the agonists, isoprenaline, adrenaline, noradrenaline, formoterol, salbutamol, salmeterol and BI-167-107, displacing the fluorescent antagonist CA200693(-S)-propranolol-green from the human β_2_AR when solubilised into a DDM micelle. These data showed all ligands bound the β_2_AR and had broad ranging p*K*_i_ values, (see **Table 1**) with the lowest affinity ligand being noradrenaline (p*K*_i_ = 4.40 ± 0.09) and highest BI-167-107 (p*K*_i_ = 9.20 ± 0.08) (n=3 ± SEM). These observed affinity values being consistent with the literature.

**Figure 1:**
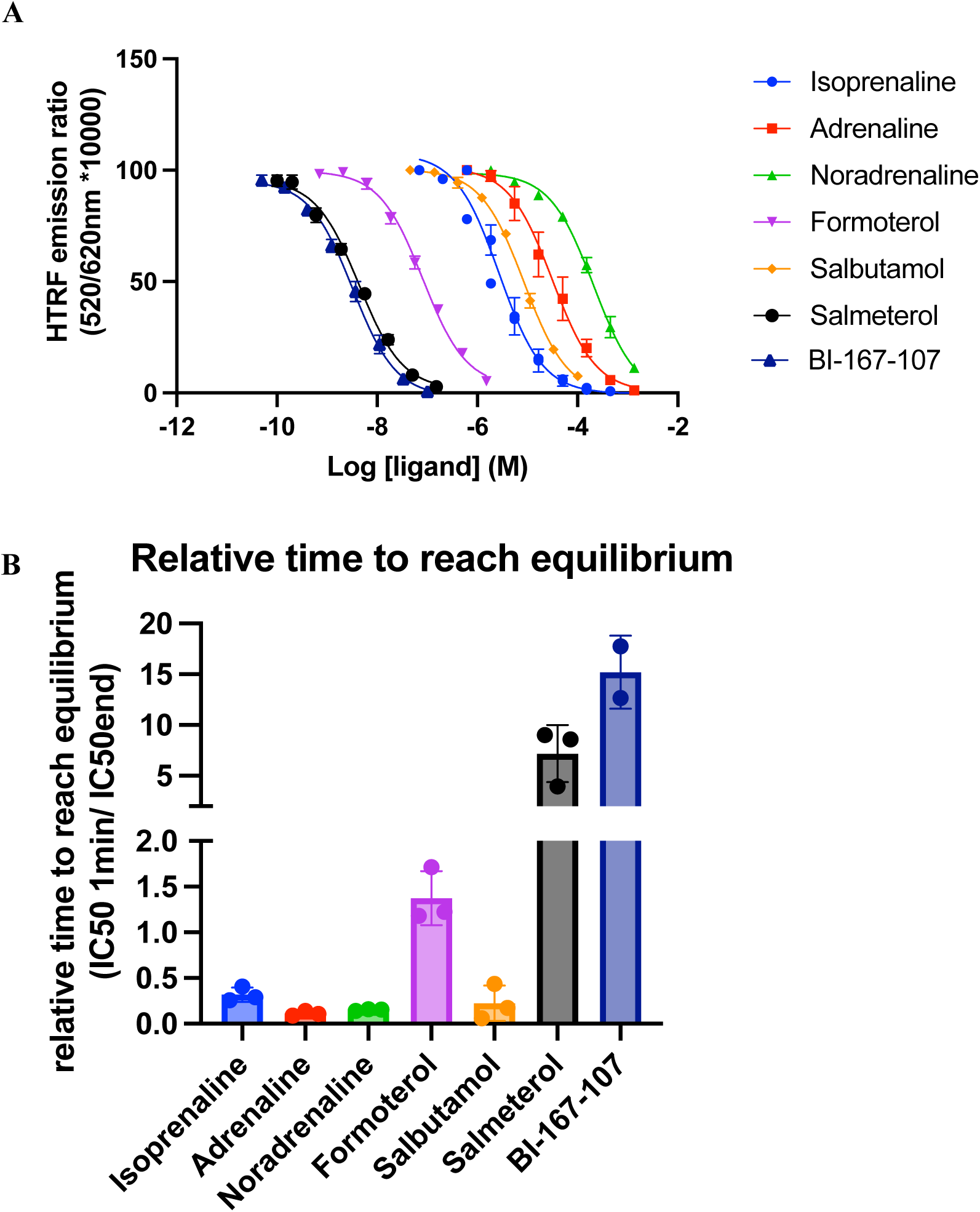
Competition binding studies for isoprenaline, adrenaline, noradrenaline, formoterol, salbutamol, salmeterol, BI-167-107 binding Lumi4-Tb labelled DDM-TS-SNAP-β_2_AR, using 75nM CA200693 (S)-propranolol-green. **A)** Equilibrium measurements were read at 20 min post DDM-TS-SNAP-β_2_AR addition for all compounds except BI-167-107 which was read at 40 min. TR-FRET between Lumi4-Tb and CA200693 (S)-propranolol-green PHERAstar FSX using two laser flashes per cycle and 520/620 TRF module. Data points show mean of three experiments normalised to 0% inhibition of specific CA200693 (S)-propranolol-green bound for each compound, ± SEM. **B)** Comparison of IC_50 1min_ / IC_50 equilibrium_ values for all seven compounds binding DDM-TS-SNAP-β_2_AR using TR-FRET, bars show mean of three experiments ± SEM.

**Table 1:** A summary of pK_i_ and IC_50 1min_ / IC_50 equilibrium_ values for isoprenaline, adrenaline, noradrenaline, formoterol, salbutamol, salmeterol, and BI-167-107 binding DDM-TS-SNAP-β_2_AR obtained from equilibrium competition binding. TR-FRET between Lumi4-Tb and CA200693 (S)-propranolol-green PHERAstar FSX using 2 laser flashes per cycle and 520/620 TRF module. Values are mean of three experiments ± SEM.

| | $pK_i \pm \text{SEM}$ | $\text{IC}_{50 \text{ 1min}} / \text{IC}_{50 \text{ equilibrium}} \pm \text{SEM}$ |
| --- | --- | --- |
| Isoprenaline | 6.4<br>$\pm 0.12$ | 0.32<br>$\pm 0.05$ |
| Adrenaline | 5.2<br>$\pm 0.25$ | 0.11<br>$\pm 0.01$ |
| Noradrenaline | 4.4<br>$\pm 0.09$ | 0.15<br>$\pm 0.01$ |
| Formoterol | 7.8<br>$\pm 0.07$ | 1.37<br>$\pm 0.17$ |
| Salbutamol | 5.8<br>$\pm 0.06$ | 0.22<br>$\pm 0.11$ |
| Salmeterol | 9.1<br>$\pm 0.02$ | 7.2<br>$\pm 1.63$ |
| BI-167-107 | 9.2<br>$\pm 0.08$ | 15.2<br>$\pm 2.08$ |

To investigate the correlation between ligand residence time and efficacy, we further characterised the ligand binding kinetics of these seven β_2_AR agonists. Whilst it is not possible to obtain a fluorescent ligand with fast enough ligand binding kinetics to accurately quantify the precise *k*_on_ and *k*_off_ rates of all of these unlabelled agonists, we were able to rank their relative time to reach equilibrium based on the difference in individual ligand IC_50_ values at 1 minute compared to IC_50_ at equilibrium (**Figure 1B**, **Table 1**). The relative time to reach equilibrium is dependent on a ligands dissociation rate, and therefore this ratio is indicative of the ligand dissociation rate constant or *k*_off_, and therefore an indication of the relative ligand residence times, with residence time being defined as 1/ *k*_off_ (Heise et al., 2007). This analysis showed a broad range of relative time to equilibrium values, with the fastest being adrenaline and noradrenaline (0.11 ± 0.01 and 0.15 ± 0.01) and the slowest being BI-167-107 (15.2 ± 2.08).

The efficacy of these seven agonists to activate the heterotrimeric G_s_ protein was then determined using a BRET-based G_s_ protein biosensor in HEK293 cells (Schihada et al., 2021). G_s_ protein activation was observed as a decrease in BRET as nLuc labelled G_sα_ dissociated from Venus tagged G_γ_ (**Figure 2A**). As expected, all compounds activated the CASE G_s_ protein (**Figure 2B-H)**, and these data were fitted to the operational model of agonism with *K*_A_ values fixed to affinity values obtained in the binding study, and the resulting log 1² values summarised in **Table 2**. These data showed a range of efficacy values with the lowest efficacy agonists being salmeterol and salbutamol and the highest being isoprenaline and adrenaline.

**Figure 2:**
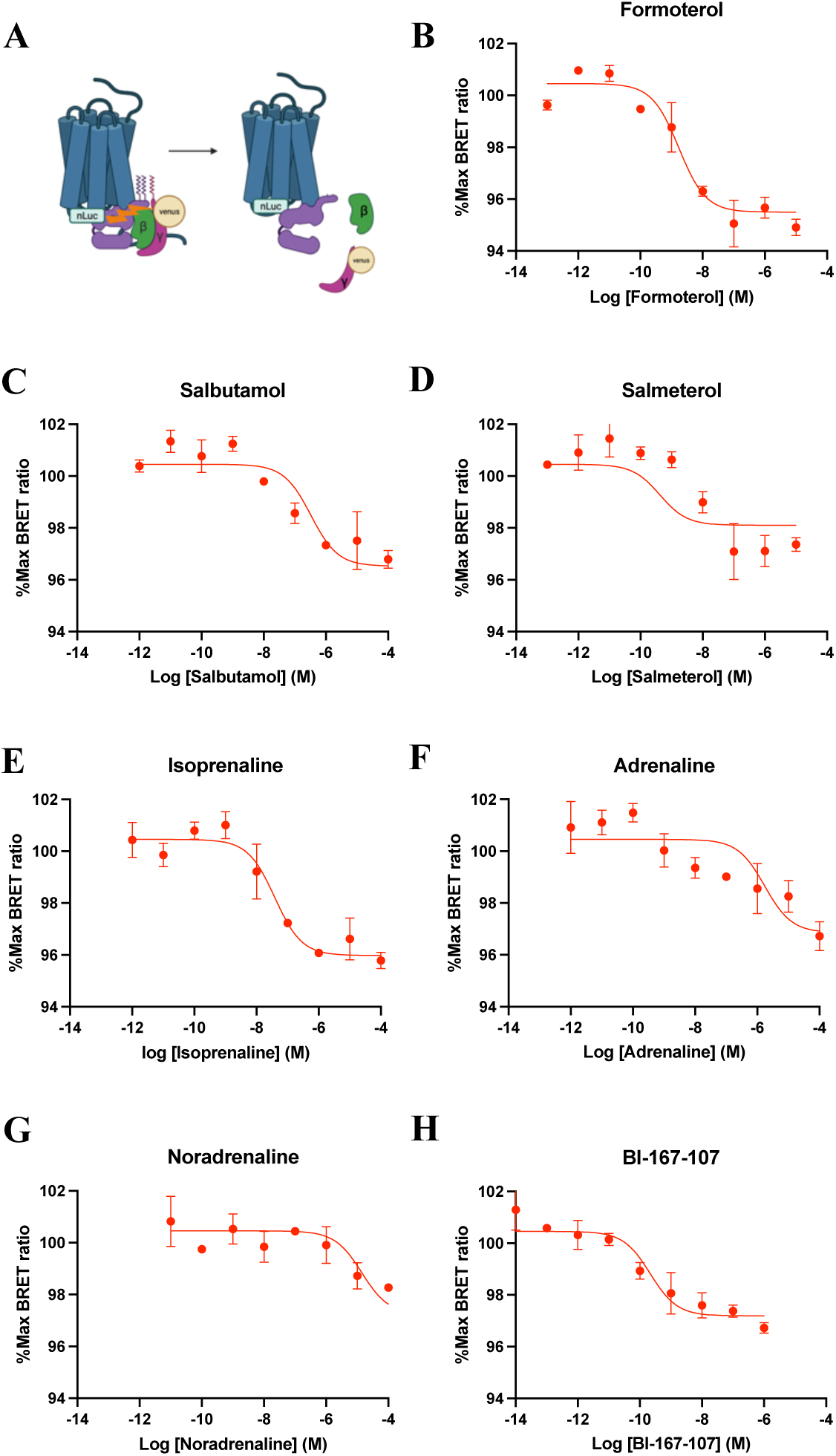
**CASE G_s_ activation studies**, summarised in **A)** for **B)** Formoterol, **C)** Salbutamol, **D)** Salmeterol, **E)** Isoprenaline, **F)** Adrenaline, **G)** Noradrenaline, and **H)** BI-167-107 were fitted to the operational model. Formoterol was used as the reference ligand and *K*_A_ values fixed to experimentally obtained *K*_i_ values. CASE G_s_ activation was obtained in a T-REx^TM^-293 -SNAP-β_2_AR and CASE G_s_ single colony population which had been induced with 1μg/mL tetracycline for 48h. Duplicate wells of adherent cells were stimulated with varying concentrations of ligand and BRET was read at 15min post ligand addition using 550LP/450-80nm luminescence module and PHERAstar FSX, Data points show mean of duplicate wells from a representative single experiment, error bars show SD.

**Table 2:** A summary of efficacy and potency values obtained for CASE-G_s_ activation by adrenaline, isoprenaline, salbutamol, BI-167-107, noradrenaline, formoterol, and salmeterol in a single colony T-REx^TM^-293 -SNAP-π_2_AR and CASE G_s_ stable cell line which had been induced with 1μg/mL tetracycline for 48h, pEC_50_ values are mean of three individually experiments individually fitted to a sigmoidal curve, E_max_ values were obtained from sigmoidal curve fits in Figure 2, log *1″* values are mean of three individually experiments individually fitted to the operational model with K_A_ values fixed to experimentally obtained K_i_ values, All error bars show SEM.

| **Ligand** | **pEC<sub>50</sub>** | **E<sub>max</sub>** | **Log $\tau$** |
| --- | --- | --- | --- |
| **Adrenaline** | 7.2 | 97.2% | 1.26 |
| | $\pm 0.5$ | | $\pm 0.84$ |
| **Isoprenaline** | 7.5 | 95.6% | 1.64 |
| | $\pm 0.23$ | | $\pm 0.96$ |
| **Salbutamol** | 7.5 | 96.8% | 0.40 |
| | $\pm 0.19$ | | $\pm 0.09$ |
| **BI-167-107** | 8.8 | 96.8% | 0.36 |
| | $\pm 0.62$ | | $\pm 0.04$ |
| **Noradrenaline** | 5.6 | 96.9% | 0.52 |
| | $\pm 0.33$ | | $\pm 0.45$ |
| **Formoterol** | 8.7 | 95.31% | - |
| | $\pm 0.18$ | | |
| **Salmeterol** | 8.0 | 97.3% | 0.05 |
| | $\pm 0.18$ | | $\pm 0.05$ |

### Lack of correlation between ligand residence time and agonist efficacy at β2AR

The correlation between agonist efficacy, as determined by log(*t*) values obtained from CASE G_s_ activation assays and ranked ligand residence time (IC_50 1min_ / IC_50 equilibrium_), obtained from TR-FRET competition ligand binding studies was determined using Pearson’s correlation analysis, see **Figure 3**. This showed no statistically significant correlation (R^2^=0.26, p=0.29) between the relative ligand residence times (IC_50 1min_ / IC_50 equilibrium_) and the efficacy values for these seven β_2_AR values.

**Figure 3:**
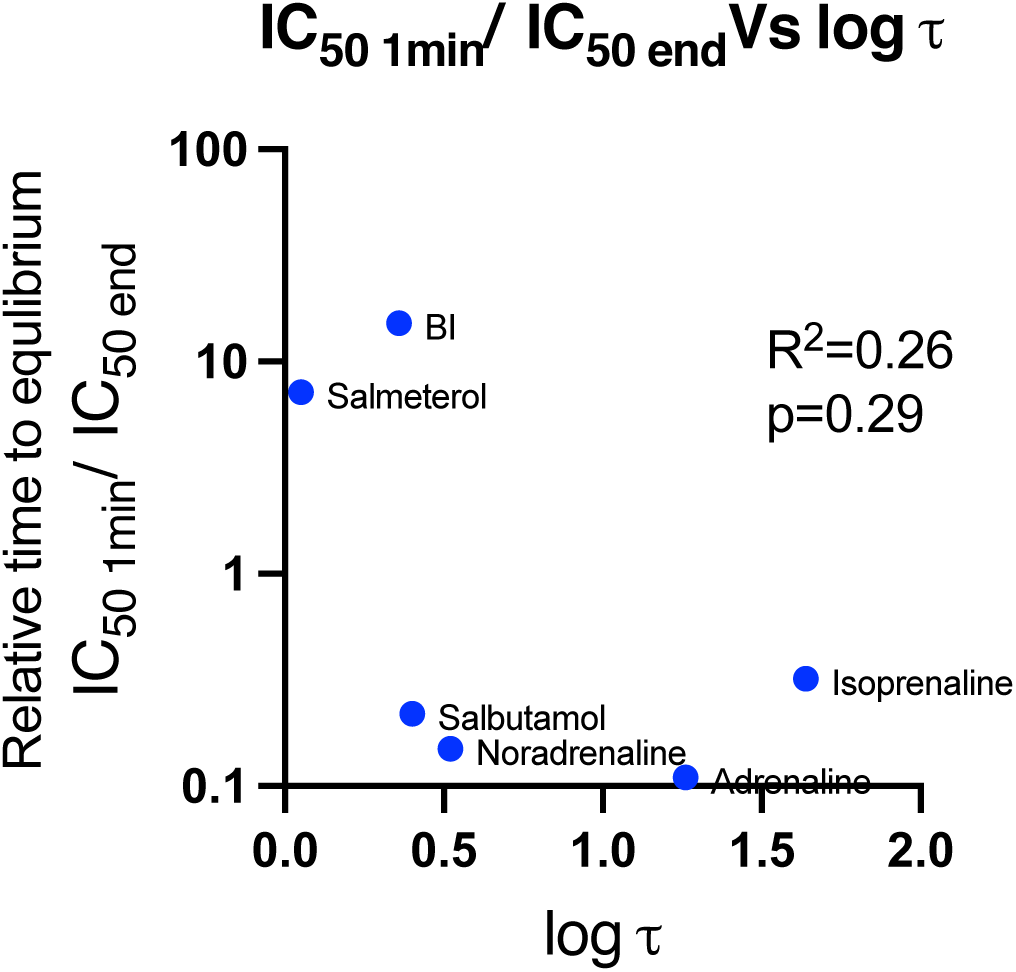
Pearsons’ correlation between ligand efficacy (log τ) values and the relative time for these ligands to reach equilibrium (IC_50 1min_ / IC_50 equilibrium_ values), data points show the mean values calculated from 3 independent experiments.

### Investigating the kinetics of mini-Gs protein binding to the β2AR in complex with agonists of varying efficacies

To probe the binding affinity and kinetics of the G_s_ protein to the agonist-β_2_AR-nLuc complex, we expressed and purified fluorescently labelled and unlabelled mini-G_s_ proteins (see **SF1**) from *E*. coli and used these in an in solution-based nanoBRET β_2_AR-nLuc binding assays.

All of the β_2_AR agonists characterised above were able to recruit the Venus-labelled mini-G_s_ protein (see **Figure 4**, **Table 3, SF2**) in a concentration-dependent fashion, and did so in rank order of their ligand binding affinities (**Table 1**), with pEC_50_ values that were similar to their p*K*_i_ values. Indeed, for the agonists adrenaline (p*K*_i_ = 5.2 ± 0.3 vs pEC_50_ =5.8 ± 0.4), noradrenaline (p*K*_i_ = 4.4 ± 0.1 vs pEC_50_ = 4.5 ± 0.3), formoterol (p*K*_i_ = 7.8 ± 0.1 vs pEC_50_ = 8.0 ± 0.2), isoprenaline (p*K*_i_ = 6.4 ± 0.1 vs pEC_50_ =7.9 ± 0.4), salbutamol (p*K*_i_ = 5.8 ± 0.1 vs pEC_50_ = 6.2 ± 0.4), and salmeterol (p*K*_i_ =9.1 ± 0.02 vs pEC_50_ = 8.7 ± 0.2), there was no difference between the pEC_50_ value obtained for Venus-mini-G_s_ recruitment and ligand binding affinity values (all p>0.05, unpaired t-test). In contrast, BI-167-107 (p*K*_i_ = 9.2±0.1 vs p*EC_50_* =8.6 ± 0.1, p=0.01) showed statistically significant differences in pEC*_50_* and p*K*_i_ values, but these values were still similar (∼3-fold different).

**Figure 4:**
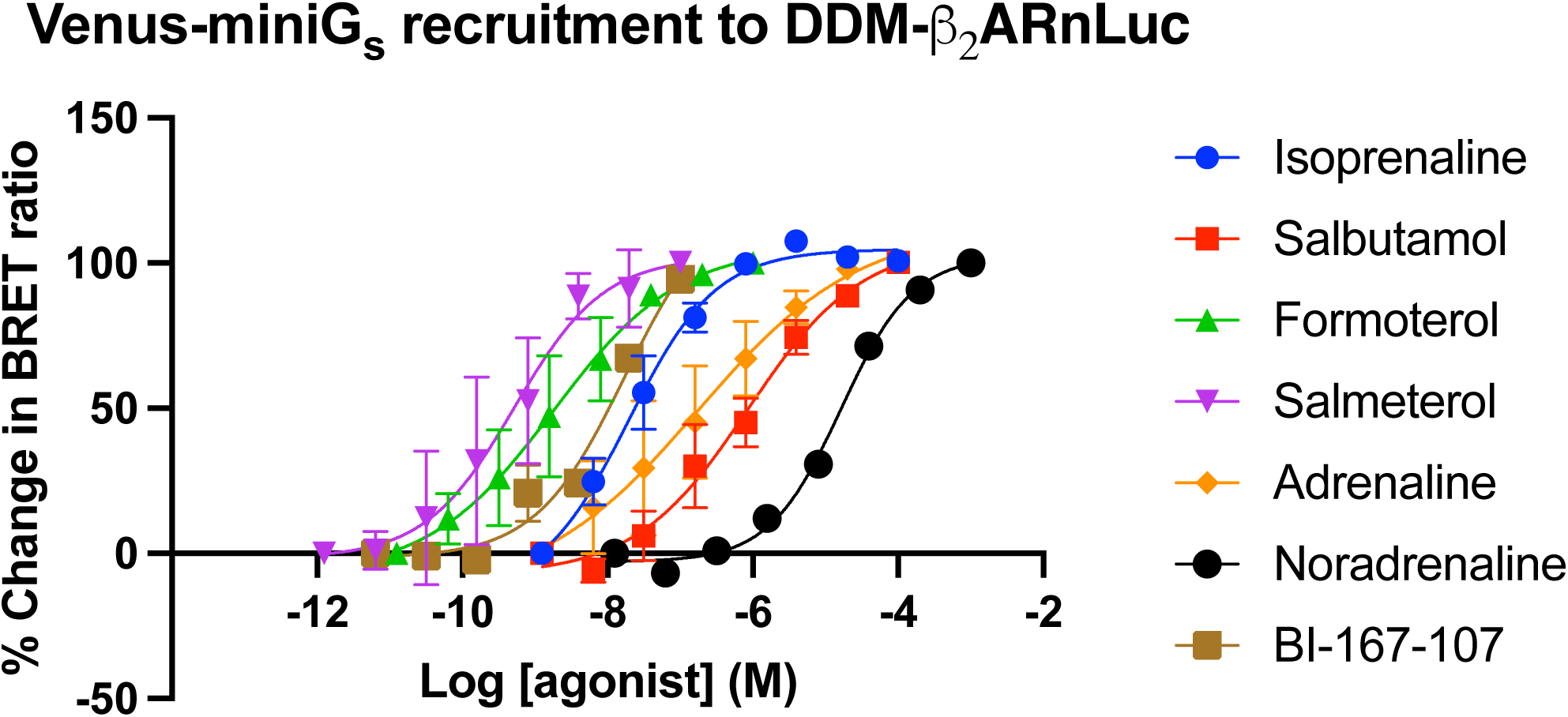
Venus-mini-G_s_ recruitment to DDM-TS-SNAP-β_2_AR-nLuc in response to different β_2_AR agonists: 1μM purified Venus-mini-G_s_ was incubated with DDM-β_2_AR-nLuc and varying concentrations of isoprenaline, salbutamol, formoterol, salmeterol, adrenaline, noradrenaline, or BI-167-107 at a final concentration 1% DMSO for 90min. NanoBRET between TS-SNAP-β_2_AR-nLuc and Venus-mini-G_s_ was read on the PHERAstar FSX, at room temperature, using LUM 550LP/450-80nm module. All curves show combined normalised mean data ± SEM of three independent observations.

**Table 3:** A summary of mean pEC_50_ values for purified Venus-mini-G_s_ recruitment to DDM-TS-SNAP-β. _2_**AR-nLuc** by various β_2_AR agonists, nanoBRET between TS-SNAP-β_2_AR-nLuc and Venus-mini-G_s_ read on PHERAstar FSX, at room temperature, using LUM 550LP/450-80nm module. pEC_50_ values show mean of n=3 individually fitted experiments ± SEM.

|  | **pEC<sub>50</sub>** |
| --- | --- |
| Isoprenaline | 7.9 ± 0.4 |
| Salbutamol | 6.2 ± 0.4 |
| Formoterol | 8.0 ± 0.2 |
| Salmeterol | 8.7 ± 0.2 |
| Adrenaline | 5.8 ± 0.4 |
| Noradrenaline | 4.5 ± 0.3 |
| BI-167107 | 8.6 ± 0.1 |

Having confirmed that all investigated agonists recruited the Venus-mini-G_s_ protein, we established a kinetic nanoBRET binding assay to measure Venus-mini-G_s_ protein recruitment to the DDM solubilised β_2_AR-nLuc. We pre-incubated the receptor with a saturating concentration (**ST1**) of each β_2_AR agonists characterised above, before adding it to a plate containing various concentrations of Venus-mini-G_s_ protein and measured the association of these two proteins using nanoBRET. The unlabelled mini-G_s_ was used to define non-specific binding and to dissociate the agonist-β_2_AR-nLuc-Venus-mini-G_s_ complex. The concentration (30μM) required for complete dissociation was calculated from mini-G_s_ competition binding studies (**SF3** and validated experimentally (**SF2, SF4**)). Hence, both association and dissociation of Venus-mini-G_s_ for the agonist-β_2_AR-nLuc could be measured (see **Figure 5**). These studies showed association of the Venus-mini-G_s_ for the agonist bound receptor to be biphasic and its dissociation to be incomplete on addition of unlabelled mini-G_s_. The biphasic association (*k*_obs_) showed a fast phase referred to as *k*_fast_ and a slow phase referred to as *k*_slow_. There was no difference in the fraction of the *k*_fast_ and *k*_slow_ of the association (*k*_obs_) across the seven β_2_AR agonists studied (**see ST2**). The average percentage of complexes dissociated was very similar across the seven agonists, ranging from 72-80% (see **ST3**), p=0.47, one-way ANOVA). This correlated with the percentage of association that was attributed to *k*_fast._

**Figure 5:**
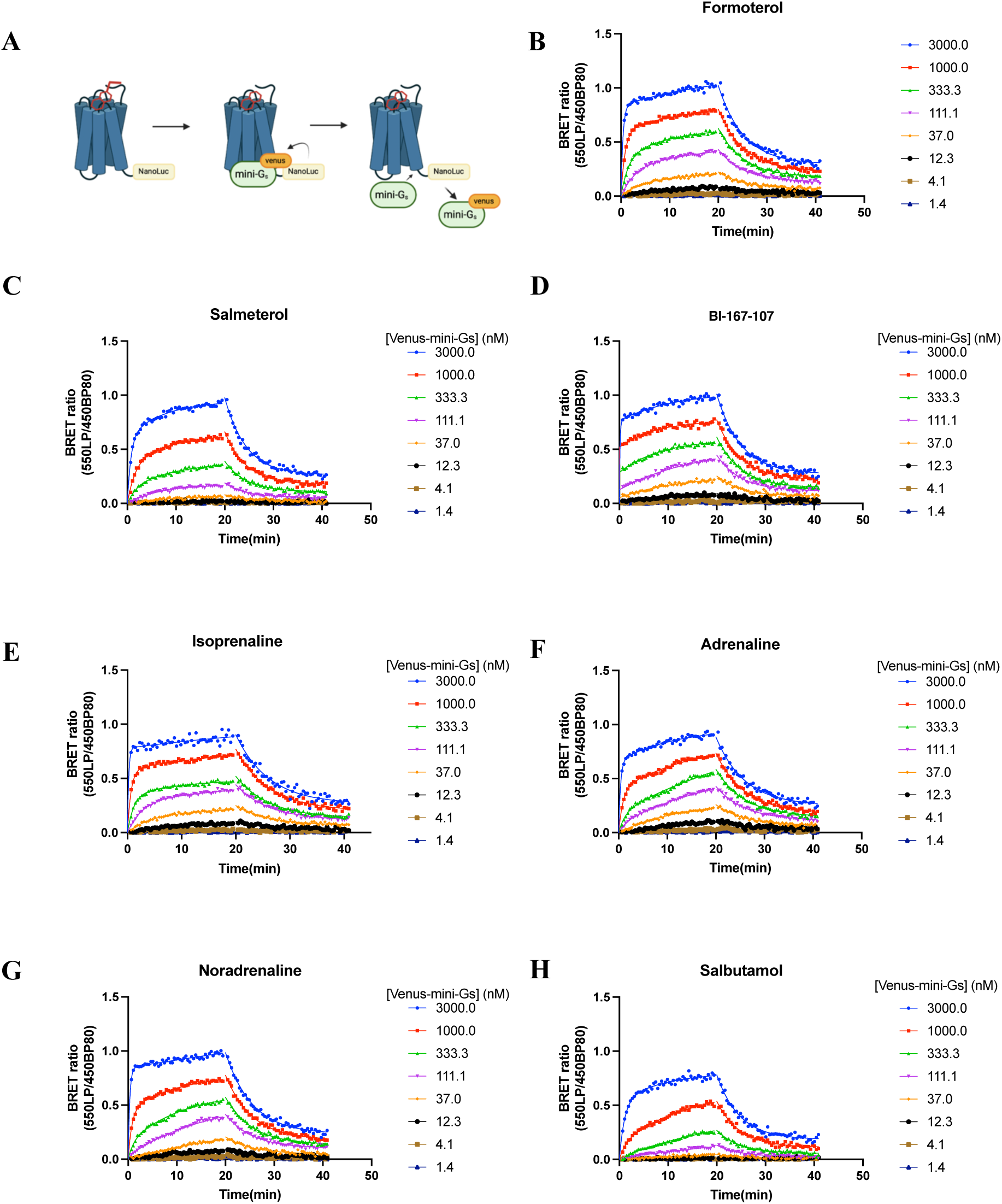
Investigation of the association and dissociation at 20min using 33μM mini-G_s,_ of Venus-mini-G_s_ binding to DDM-TS-SNAP-β_2_AR when preincubated with saturating concentration of B) Formoterol C) Salmeterol D) BI-167-107 E) Isoprenaline F) adrenaline G) Noradrenaline H) Salbutamol at room temperature, using LUM 550LP/450-80nm module. All figures show specific binding, where 30μM mini-G_s_ was used to define the NSB, representative raw data of n=3, fitted to a two-phase association and one phase dissociation.

Whilst *k*_obs_ plots for *k*_slow_ did not follow a linear relationship (**SF5**), *k*_obs_ plots for *k*_fast_ followed a linear relationship (**SF6**) for the majority of the ligands and were used to calculate the *k*_on_ of *k*_fast_ (*k*_on(fast)_), which appeared the most relevant parameter for considering G protein recruitment in our study (see **Table 4**). These *k*_on(fast)_ values for Venus-mini-G_s_ binding agonist bound DDM-TS-SNAP-β_2_AR-nLuc were in the range of 3.4 ± 0.64 to 9.2 ± 0.26 x10^5^ M^-1^ Min ^-1^ (see **Table 4**). A one-way ANOVA showed that this variation in range was statistically significant (p=0.03), but Tukey’s multiple comparison test shown no statistically significant difference in pairwise comparisons. There was very little variation in *k*_off_ values for the Venus-mini-G_s_ dissociating from the agonist bound DDM-TS-SNAP-β_2_AR-nLuc, showing a range of 0.17 to 0.21 min^-1^. Subsequently residence times for the Venus-mini-G_s_ were all approximately 5 minutes (see **Table 4**).

**Table 4:** A summary of the mean pK_d_, k_off_, k_on(fast)_ and residence time values for Venus-mini-G_s_ proteins binding to the DDM solubilised TS-SNAP-β_2_AR bound to the β_2_AR agonists BI-167107, formoterol, isoprenaline, adrenaline, noradrenaline, salbutamol and salmeterol, as measured by nanoBRET, values show mean of n=3-4 experiments ± SEM. ^%^denotes full agonist and ^$^partial agonist, as defined by CASE G_s_ activation assay.

|  | **pK<sub>d</sub>** | **K<sub>off</sub>**<br>**(Min<sup>-1</sup>)** | **K<sub>on(fast)</sub>**<br>**(M<sup>-1</sup> Min<sup>-1</sup>)** | **Residence time**<br>**(Min)** |
| --- | --- | --- | --- | --- |
| %BI-167107 | 6.7<br>±0.03 | 0.21<br>±0.003 | 7.29 ± 2.16<br>x10<sup>5</sup> | 4.76 |
| %Formoterol | 6.7<br>±0.05 | 0.20<br>±0.011 | 4.59± 1.64<br>x10<sup>5</sup> | 5.00 |
| %Isoprenaline | 6.8<br>±0.07 | 0.17<br>±0.004 | 9.19± 0.42<br>x10<sup>5</sup> | 4.76 |
| %Adrenaline | 6.8<br>±0.05 | 0.18<br>±0.014 | 8.56± 0.13<br>x10<sup>5</sup> | 5.50 |
| %Noradrenaline | 6.6<br>±0.08 | 0.19<br>±0.007 | 7.92 ±0.56<br>x10<sup>5</sup> | 5.20 |
| \$Salbutamol | 6.0<br>±0.07 | 0.20<br>±0.006 | 3.36±0.64<br>x10<sup>5</sup> | 5.00 |
| \$Salmeterol | 6.1<br>±0.09 | 0.21<br>±0.006 | 4.18±1.2<br>x10<sup>5</sup> | 4.76 |

Affinity p*K*_d_ values for Venus-mini-G_s_ binding the agonist:DDM-TS-SNAP-β_2_AR-nLuc complex were obtained by fitting association data at 20 min to a one-site saturation specific binding model as shown in **Figure 6**. Overall, these data show small differences in p*K*_d_ values that were driven by small increases in *k*_on(fast)_ values. There were 0.5-0.8 log units of increased affinity for Venus-mini-G_s_ binding the full agonist (adrenaline, noradrenaline, formoterol, isoprenaline, and BI-167-107) bound DDM-TS-SNAP-β_2_AR-nLuc complex compared to the partial agonists (salbutamol and salmeterol) bound, which proved a statistically significant difference (one-way ANOVA and Tukey’s post hoc).

**Figure 6:**
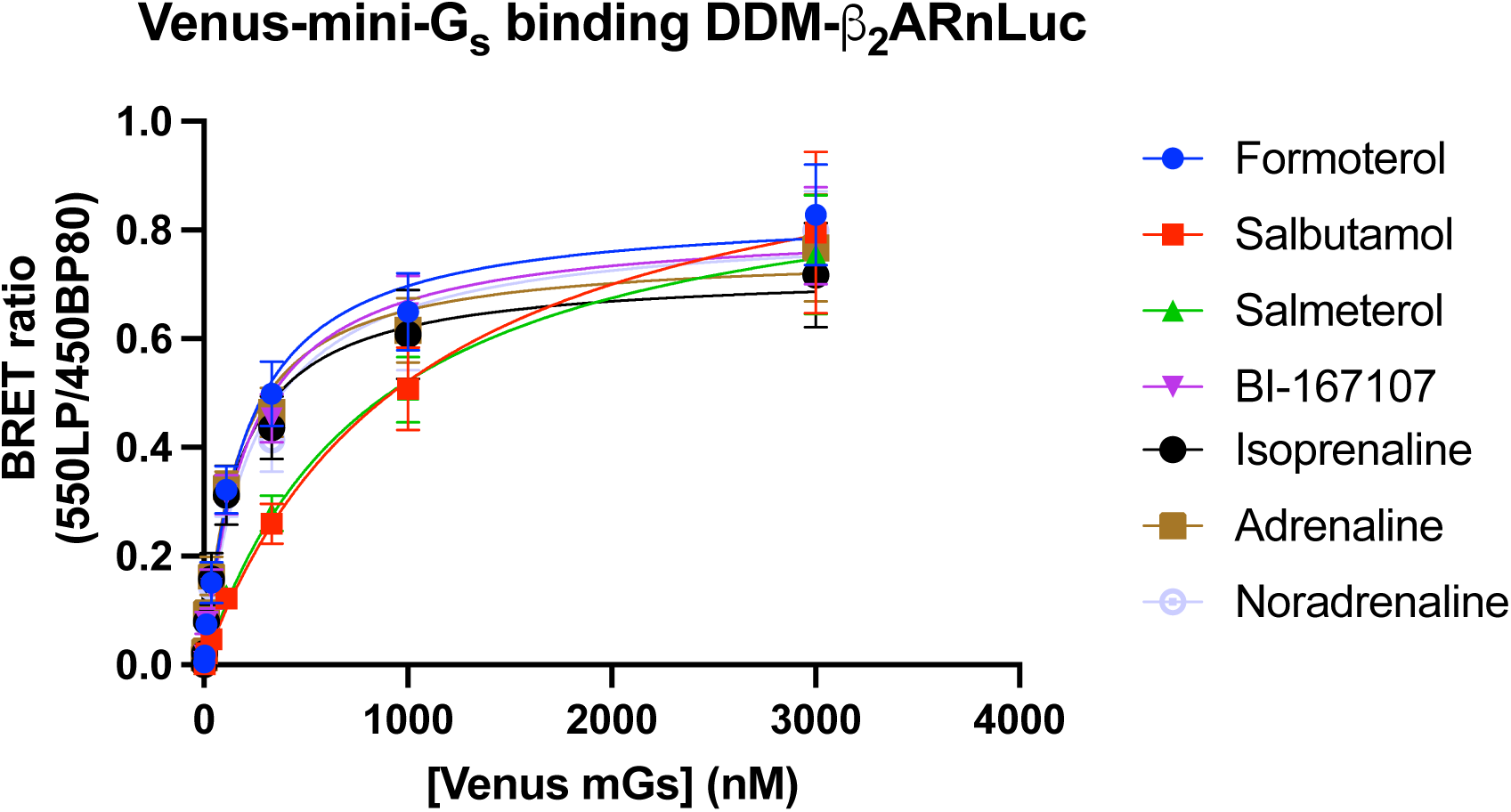
Specific saturation binding of increasing concentrations of purified Venus-mini-G_s_ binding to DDM-TS-SNAP-β_2_AR-nLuc in the presence of saturating concentrations of formoterol, salbutamol, salmeterol BI-167-107, isoprenaline, adrenaline and noradrenaline. nanoBRET between TS-SNAP-β_2_AR-nLuc and Venus-mini-Gs was read on PHERAstar FSX, at room temperature, using LUM 550LP/450-80nm module at 20min. Data was fitted to one-site specific binding model, points show the mean ± SEM of 4 independent experiments.

Overall, there were significant differences in the affinity of the Venus-mini-G_s_ complex for the DDM-TS-SNAP-β_2_AR-nLuc, when in complex with full β_2_AR agonists compared to partial β_2_AR agonists, and this difference was driven by an increased *k*_on(fast)_, with no difference in *k*_off_.

### Affinity and the rate of association of Venus-mini-Gs protein for β2AR-nLuc correlated with agonist efficacy

Finally, we performed Pearson’s correlation analysis between both the association rate (*k*_on(fast)_) and affinity (p*K*_d_) values for Venus-mini-G_s_ binding agonist-β_2_AR-nLuc complexes vs agonist efficacy or log τ, a measure of the ability of each agonist to activate the heterotrimeric G_s_ protein values (see **Figure 7**). This analysis showed a moderate correlation between both ligand efficacy (log *1″*) and mini-G_s_ association rate (*k*_on(fast)_) (R^2^=0.58, p=0.07) and between ligand efficacy (log *1″*) and mini-G_s_ affinity (R^2^=0.50, p=0.11). This suggests that the differences in agonist efficacy can be explained by agonist-β_2_AR complexes ability to recruit the G_s_ protein.

**Figure 7:**
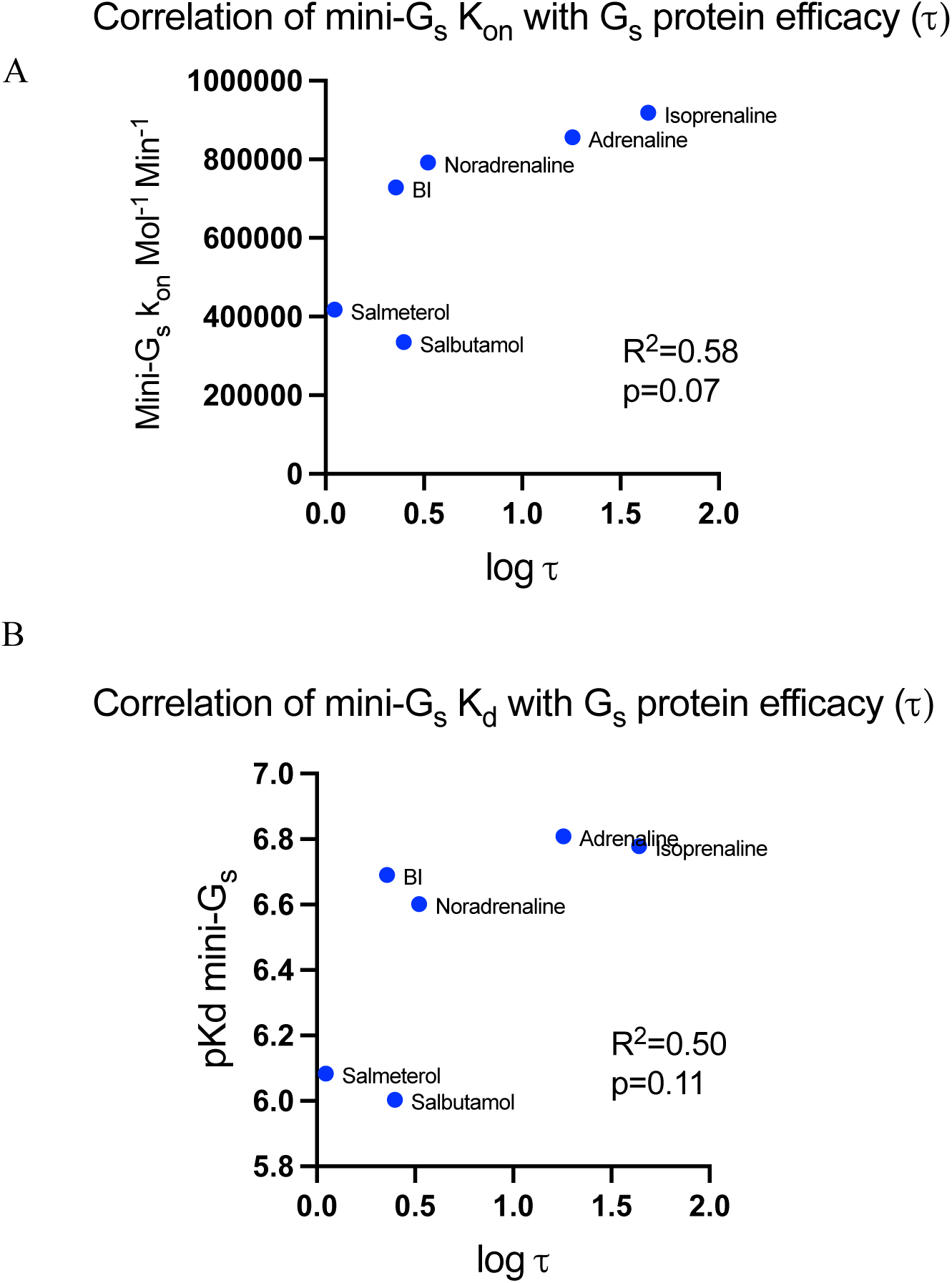
The efficacy of ligands correlates with with G protein recruitment k_on_. Correlation of CASE G_s_ activation efficacy t with A) mini-G_s_ p*K*_d_ and B) mini-G_s_ *k*_on_(_fast_), t values were obtained from fitting the operational model on individual data sets with fixed K_A_ values, correlations were made using Pearson’s correlation coefficient, on replicates from n=3-4 experiments for p*K*_d_, *k*_on_(_fast_) and *t* values.

## Discussion

Despite the therapeutic importance of GPCRs, understanding of the molecular basis of drug efficacy at GPCRs remains poorly understood. Whilst some studies suggest a role for ligand binding dissociation kinetics (Guo et al., 2012; Sykes et al., 2014) or G protein activation kinetics studies (Nikolaev et al., 2006; Gregorio et al., 2017) other studies suggest that variations in the receptor conformations induced by the agonist underly efficacy (Liu et al., 2012). We investigated the binding kinetics of seven clinically relevant agonists and the activation of mini-G_s_ at the β_2_AR and found no correlations between these parameters and efficacy.

Using the relative time to equilibrium method of Heise and colleagues (Heise et al., 2007) to obtain a relative measure of *k*_off_ and therefore a relative measure of ligand residence time (1/ IC_50 1min_/ IC_50 equilibrium_). The affinity (p*K*_i_) and rank order relative time to equilibrium (IC_50 1min_/ IC_50 equilibrium_) values for the agonists in this study generally matched those obtained by Sykes and colleagues (Sykes et al., 2014). Moreover, we used the CASE G_s_ (Schihada et al., 2021)and operational model of agonism (Black and Leff, 1983) to quantify the efficacy (log 1″) of these seven agonists and assess their ability to activate the heterotrimeric G_s_ protein. These log τ values gave a rank order of efficacy of isoprenaline > adrenaline > noradrenaline > salbutamol >BI-167-107 >salmeterol, which was similar to that of isoprenaline > adrenaline > BI-167-107 > salbutamol > salmeterol obtained by Gregorio and colleagues (Gregorio et al., 2017) who define efficacy by the effectiveness to generate G_s_ (GTP) from G_s_ (GDP).

To investigate if a longer ligand residence time correlated with efficacy at the β_2_AR we investigated if there was a correlation between relative time to equilibrium (IC_50 1min_/ IC_50 equilibrium_) and efficacy (log τ) values. We found that there was no correlation between these parameters (R^2^=0.26, p=0.29), suggesting that residence time of these β_2_AR agonists is not important in efficacy, consistent with previous studies (Sykes and Charlton, 2012). This finding contrasts with the positive correlation shown between the efficacy of agonists at the muscarinic M3 receptor and the adenosine A_2A_ receptor and their ligand residence time (Sykes et al., 2009; Guo et al., 2012). However, similarly, no correlation between efficacy and residency time was shown for the adenosine A_1_ receptor (Louvel et al., 2014). Overall, this suggests ligand residence time could be an important mechanism in the efficacy of some but certainly not all agonists binding Class A GPCRs.

Finally, we investigated the kinetics and affinity of Venus-mini-G_s_ binding to the DDM-TS-SNAP-β_2_AR-nLuc when bound to the seven β_2_AR agonists (see **Table 4**). Whilst these data showed no difference in the dissociation rate (*k*_off_) or corresponding residence time of the Venus-mini-G_s_ for the receptor when bound to the different agonists, there were statistically significant differences in the affinity (p*K*_d_) of the Venus-mini-G_s_ for the full agonist bound receptor compared to the partial agonist bound receptor. These differences appeared to be driven by an increase in the rate of association (*k*_on(fast)_). Comparison of these association rates and affinity (p*K*_d_) values for Venus-mini-G_s_ binding agonist:β_2_AR complexes with agonist efficacy values (log 1″) (see **Figure 7**). Figure 7A showed a moderate correlation between ligand efficacy (log 1″) and mini-G_s_ association rate (*k*_on(fast)_) (R^2^=0.58, p=0.07). Figure 7B shows a moderate correlation between ligand efficacy (τ) and mini-G_s_ affinity (p*K*_d_) for the agonist-β_2_AR complex (R^2^=0.50, p=0.11). These differences in the initial rate of mini-G_s_ recruitment and the resulting differences in mini-G_s_ affinity, suggest that subtle differences in agonist-β_2_AR complex conformations result in differences in agonist efficacy because of differences in the ability of these conformations to affect recruitment of Venus-mini-G_s._

This data does not, therefore, suggest a role for G protein dissociation kinetics in the molecular basis of efficacy at the β_2_AR, but suggests a model whereby full agonists stabilise a conformation of receptors which is more likely to recruit the G_s_ protein, but once bound to the receptor there is no conformational difference in the agonist-DDM-TS-SNAP-β_2_AR-nLuc-Venus-mini-G_s_ complex (**Figure 8**).These data therefore provide no evidence for a role of ligand or G protein dissociation kinetics in the molecular basis of efficacy.

**Figure 8:**
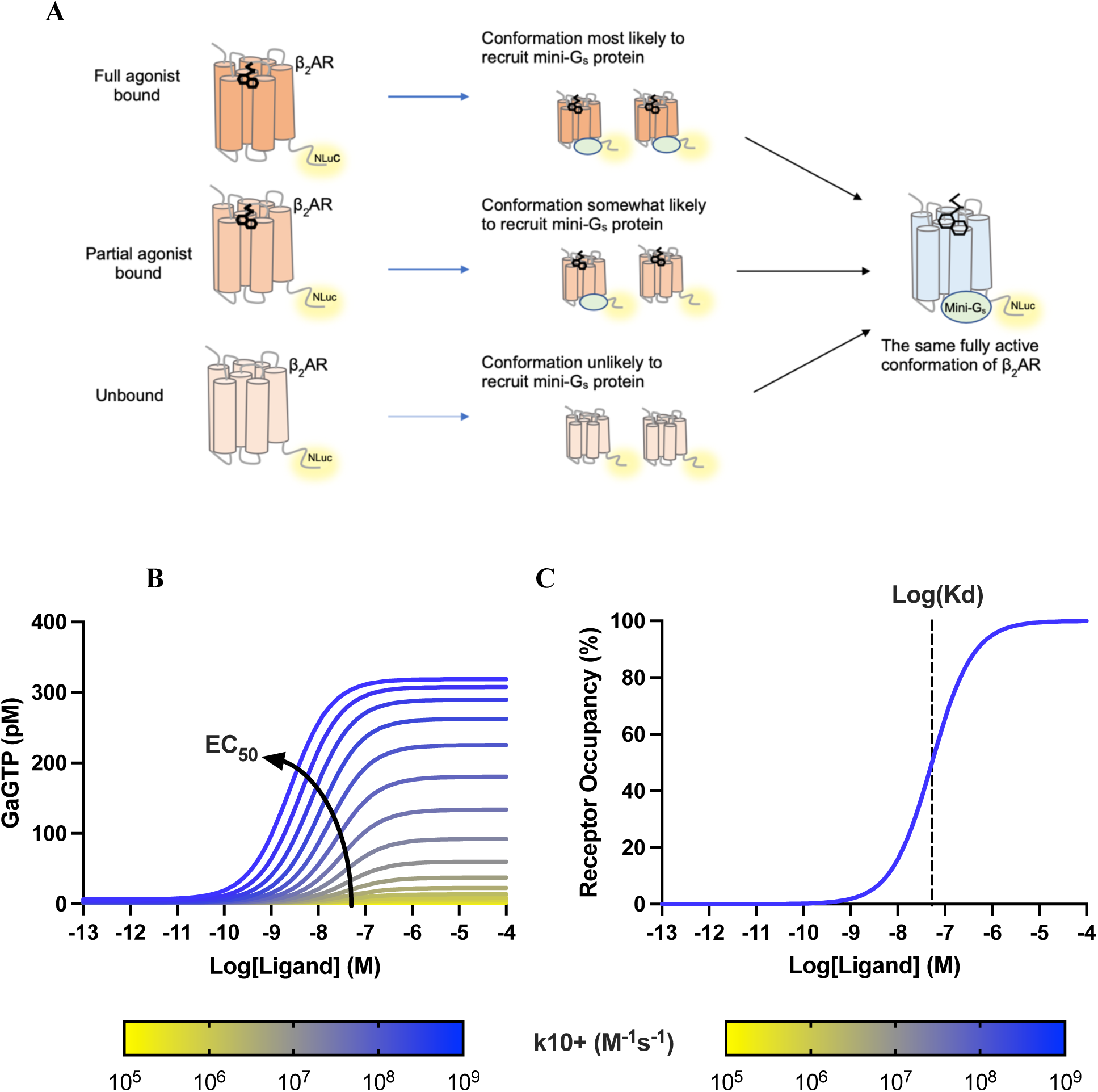
Conformational model of efficacy proposed by this study: **A)** agonists of higher efficacy induce a conformation of β_2_AR that is more likely to recruit a mini-Gs protein but once bound there is no difference in the β_2_AR conformation within the agonist-β_2_AR-mini-Gs complex, **B)** Use of the cubic ternary complex model to investigate the effect of increasing the rate of G protein recruitment on the potency of the agonist-receptor complex to activate the G protein. Arrow indicates increases in apparent ligand EC_50_ values for the formation of GαGTP. **C)** Use of the cubic ternary complex model to investigate the effect of increasing the rate of G protein recruitment on agonist affinity for the GPCR. Dotted line indicates log(*K*_d_) of ligand-receptor occupancy. As shown in the figure the association rate of Ga to the receptor does not affect ligand binding affinity, hence the yellow and blue curves lie directly on top of each other.

To support our hypothesis, we applied a previously validated mathematical model of the cubic ternary complex model (Biomodels ID:2306220001), (**SF7**, ST4), to investigate the effect of increasing the forward rate of G protein binding to activated receptor on both G protein activation and agonist-receptor occupancy of the receptor (**Figure 8B-C**). As indicated, increasing the on rate for G protein recruitment increases the efficacy and potency of G protein activation by the ligand, without changing agonist-receptor occupancy (**Figure 8B**). This therefore supports our hypothesis that an increase in likelihood of G protein recruitment underlies the molecular basis of ligand efficacy at the β_2_AR.

Moreover, this conformational model (see **Figure 8**) is supported by data from hydrogen/deuterium exchange mass spectrometry (HDMS) and hydroxy radical foot printing mass spectrometry (HDX) (Du et al., 2019), whereby the conformational changes involved in β_2_AR-G_s_ protein complex formation were investigated. Du et al showed that the conformation of the initial β_2_AR-G_s_ structure differs from that of the fully formed nucleotide free β_2_AR-G_s_ complex. Furthermore, nuclear magnetic resonance (NMR) studies (Nygaard et al., 2013; Manglik et al., 2015), show that the agonist BI-167-107 alone is not enough to fully stabilise the β_2_AR in the active state and that nanobody 80 is required to fully stabilise the active state. These data support our findings that the conformation of the agonist-β_2_AR complex differs from that of the agonist-β_2_AR-mini-G_s_ complex. Moreover, Liu and colleagues (Liu et al., 2012) investigated the conformational states of β_2_AR bound to agonists of a range of efficacies and show efficacy-dependent differences in the agonist-β_2_AR conformational state. Structural studies of the agonist bound β_2_AR or other class A GPCRs have only been possible in the presence of a G protein mimetics (Rasmussen et al., 2011), and show only very small conformational differences that do not seem to explain differences in efficacy (Katritch et al., 2009). This further supports our finding that there was no difference in the agonist-β_2_AR-mini-G_s_ conformation.

It is important to acknowledge, that in order to investigate ligand and mini-G_s_ binding kinetics to the β_2_AR purely at the biophysical level, and in isolation from regulatory elements of the cell, we studied its function in the DDM micelle. Therefore, the applicability of our findings with regard to the native cell environment remains to be elucidated. Interestingly, Sungkaworn and colleagues (Sungkaworn et al., 2017) investigated the association rate (*k*_on_) and dissociation rate (*k*_off_) of Gα_I_ binding to the α2A receptor in CHO cells in response to a range of agonists using single molecule microscopy and show efficacy is at least partially correlated with *k*_on_ but not *k*_off_ of the Gα_I._ Taken together with evidence from the current study, this suggests that the conformational model of efficacy proposed herein may translate to the cell environment. Further work would investigate if the model of efficacy proposed is relevant to the β_2_AR in its native cellular environment and would ascertain if this model is a general mechanism for agonist efficacy at other GPCRs.

## Conclusion

Overall, we find no evidence for ligand or G protein binding dissociation kinetics in the molecular basis of ligand efficacy at the β_2_AR, at the biophysical level, but suggest a conformational model of efficacy whereby agonists of higher efficacy stabilise a conformation of β_2_AR that is more likely to recruit the G protein. Further studies with a wider range of agonists of differing efficacy and measurements with agonists at other receptors would ascertain if this is a general mechanism for efficacy at GPCRs.

## Acknowledgements

C.R.H was funded by a Medical Research Council (MRC) IMPACT PhD studentship. T.R.N. was funded by a UK Biotechnology and Biological Sciences Research Council iCase studentship (BB/V509334/1) co-funded with AstraZeneca. G.L. is a Royal Society Industry Fellow (NF\R2\212001). We would like to thank Dr Bradley Hoare and Dr Franziska M. Heydenreich for respectively generating the pcDNA4TO-TS-SNAP-β_2_AR-nLuc and pcDNA4TO-TS-SNAP-β_2_AR plasmids used in this study.

## Author contributions

C.R.H. designed and performed the experiments. T.R.N. performed the computational modelling.

C.R.H. wrote the manuscript and it was edited by D.A.S., S.J.B., T.R.N., G.L. and D.B.V. D.A.S. gave technical advice.

