## supplementary data file for "Agonist efficacy at the β_2_AR is driven by agonist-induced differences in receptor affinity for the G_s_ protein, not ligand binding kinetics"

**Supplementary information**

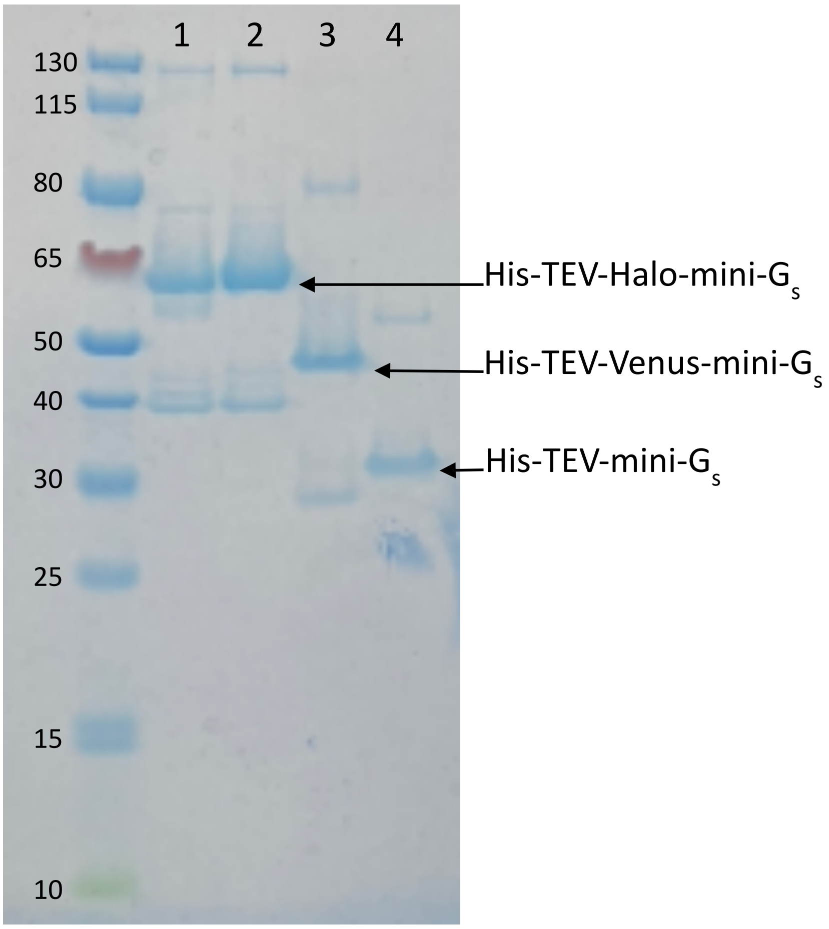


**SF1: Production of mini-$G_S$ proteins from *E.* coli:** SDS-PAGE gel stained for protein with InstantBlue showing production and purity of His10-Halo-mini-$G_s$ (lanes 1-2), His10-venus-mini-$G_s$ (lane 3) and His10-mini-$G_s$ proteins (lane 4). Representative gel of n=4 protein preps for His10-Halo-mini-$G_S$ and His10-mini-$G_S$ proteins and n=1 for His10-Venus- mini-$G_s$.

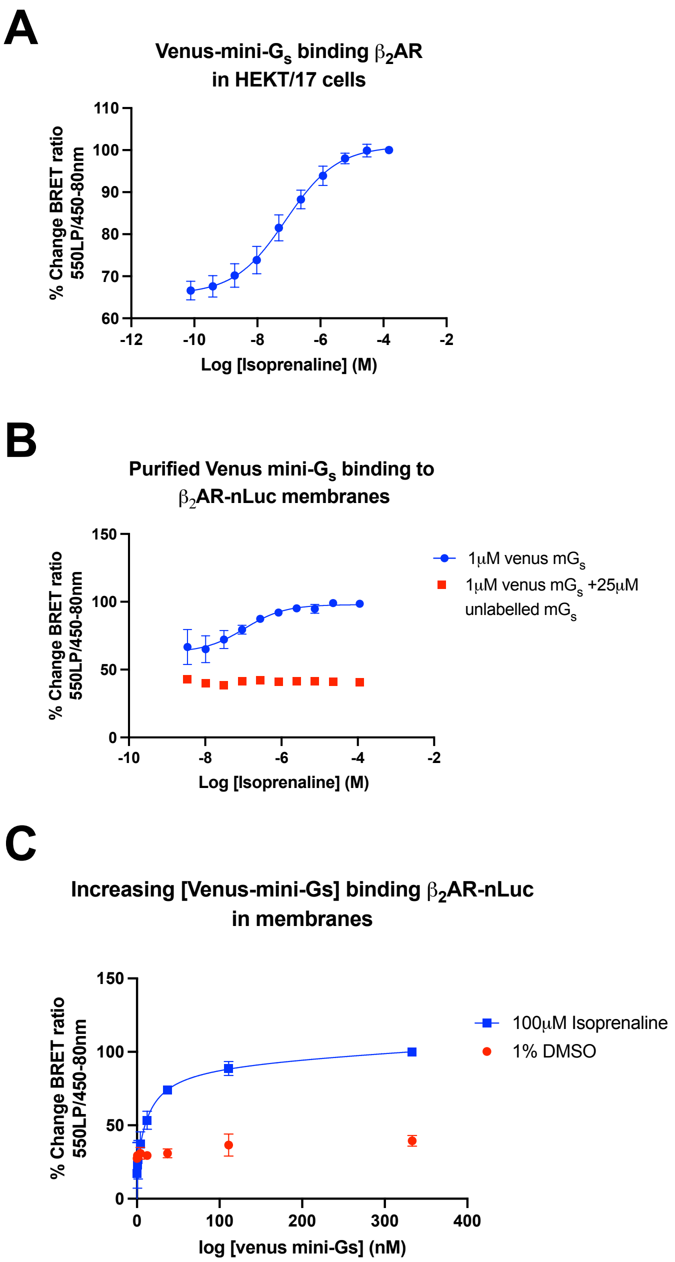


**SF2: Venus and unlabelled mini-$G_s$ proteins produced bind TS-SNAP-$\beta_2$ARnLuc. A)** Recruitment of Venus-mini-$G_s$ to TS-SNAP-$\beta_2$AR-nLuc in transiently transfected HEK293T/17 cells in response to varying concentrations of isoprenaline **B)** Recruitment of purified Venus mini-$G_s$ to membranes expressing TS-SNAP-$\beta_2$AR-nLuc in response to varying concentrations of isoprenaline **C)** Saturation binding curves for varying concentrations of purified Venus-mini-$G_s$ binding to TS-SNAP- $\beta_2$AR-nLuc membranes in the absence and presence of 100μM isoprenaline. nanoBRET between TS-SNAP-$\beta_2$AR-nLuc and Venus-mini-$G_s$ was read on PHERAstar FSX using LUM 550LP/450-80nm module. All curves show combined normalised data of n=3, error bars show ±SEM.

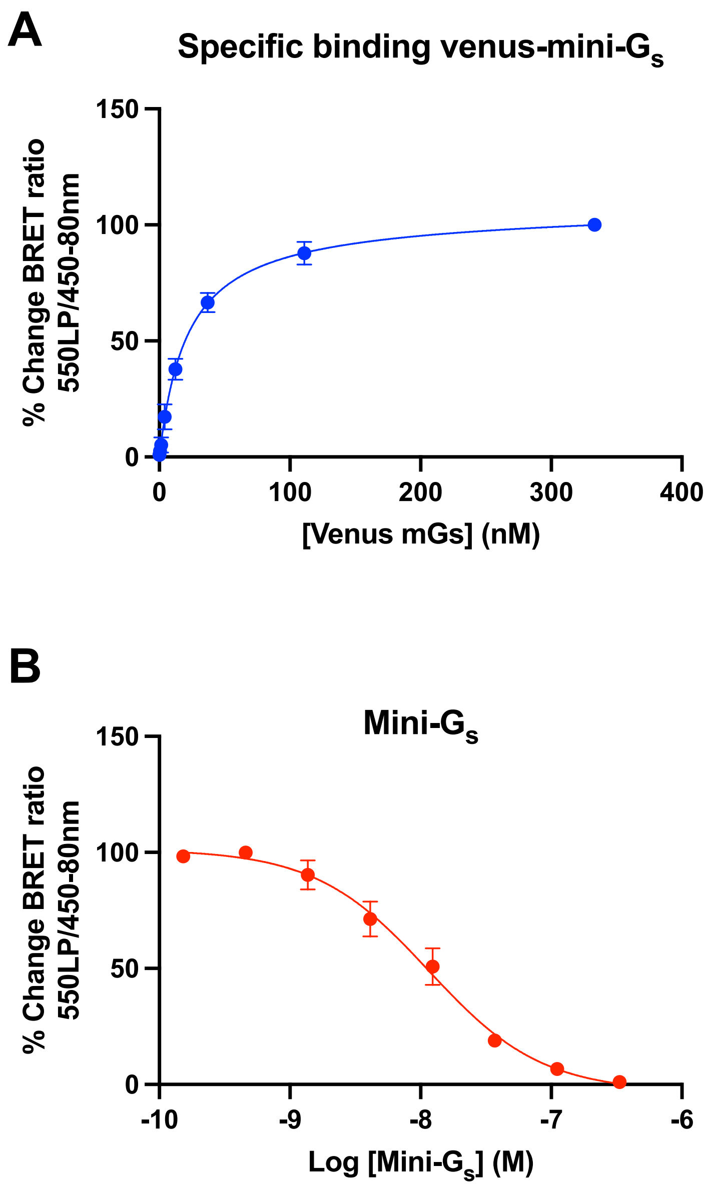


**SF3: Characterising the affinity of mini-Gs proteins for the isoprenaline bound TS-SNAP-β$_2$AR-nLuc in membranes A)** Specific saturation binding for purified Venus-mini-Gs, data points show mean±SEM, from 3 independent experiments **B)** Competition binding between 30 nM purified Venus-mini- Gs and varying concentrations of purified mini-Gs, data points show n=3± SEM, data points show n=3± SEM. BRET between TS-SNAP-**β$_2$AR-nLuc** and Venus-mini- Gs was read on PHERAstar FSX using LUM 550LP/450-80nm module. p*K*d values are mean of n=2±SD or 3±SEM individual experiments.

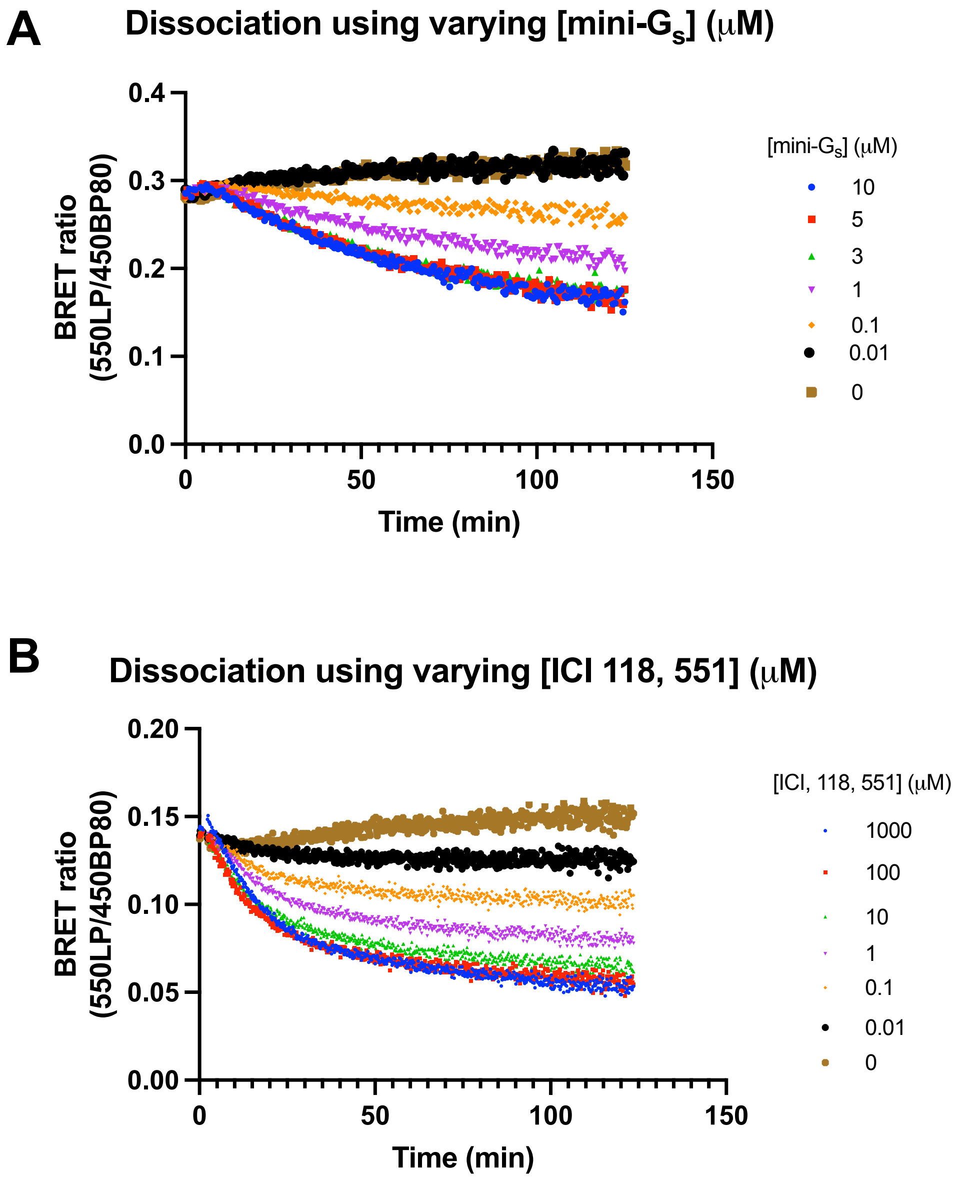


**SF4: Validation of the concentration of disociator required to dissociate the membrane-TS-SNAP-$\beta_2$AR-nLuc complex:** membrane-TS-SNAP-$\beta_2$AR-nLuc was preincubated with saturating concentration of isoprenaline and 333nM Venus-mini-$G_S$ before nanoBRET between TS-SNAP-$\beta_2$AR-nLuc and Venus-mini-$G_S$ read on PHERAstar FSX, at room temperature, using LUM 550LP/450-80nm module for 3 mins before addition of varying concentrations of **A)** mini- $G_S$ or **B)** the inverse agonist ICI 118, 551 to dissociate the membrane- SNAP-$\beta_2$AR-nLuc complex, and read for a further 2hrs, specific binding data, where unlabelled mini-$G_S$ was used to define NSB, representative data of n=3.

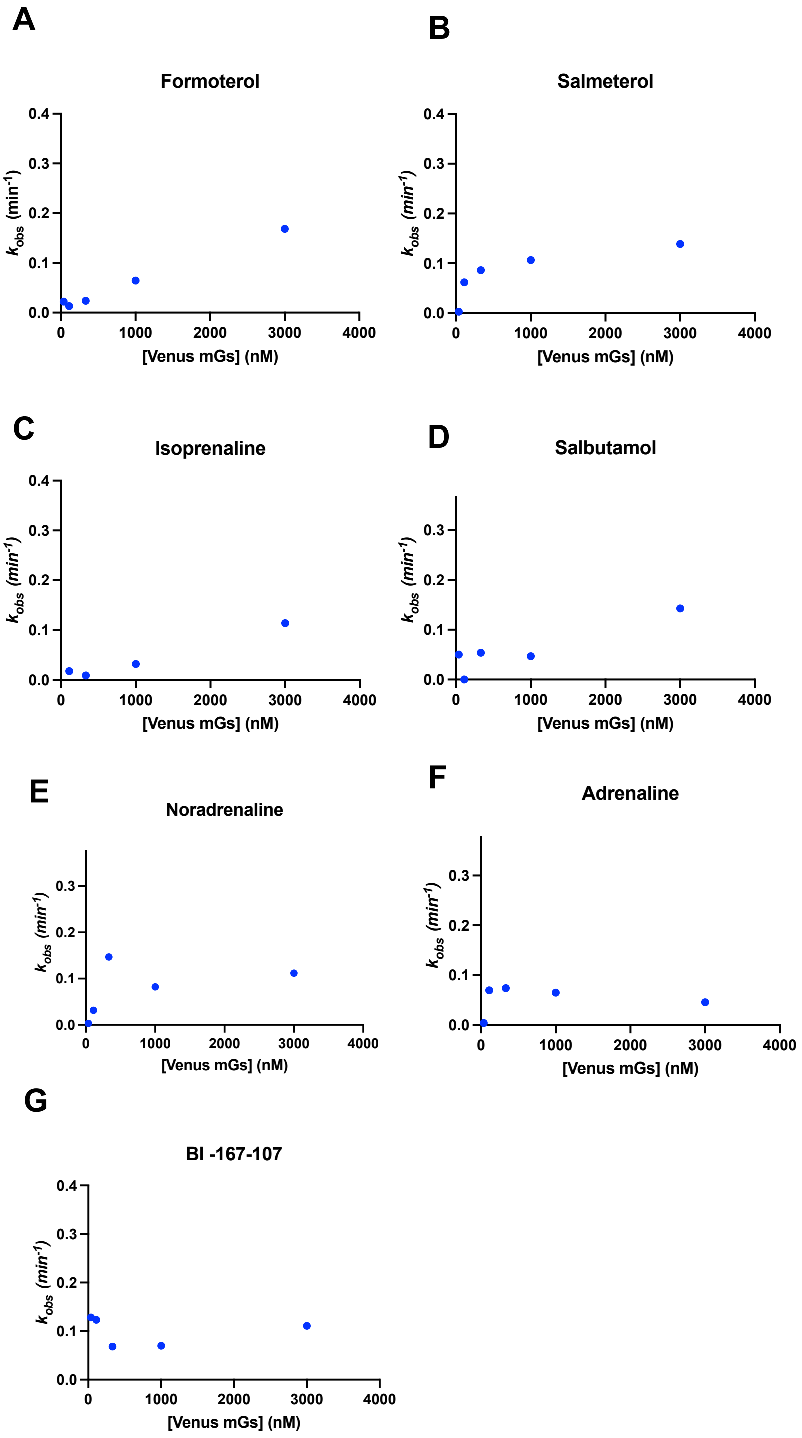


**SF5: $k_{obs}$ plots of $k_{slow}$ for Venus-mini-Gs association to DDM solubilised** TS-SNAP-$\beta_2$AR-nLuc **bound to A)** Formoterol **B)** Salmeterol **C)** Isoprenaline **D)** Salbutamol **E)** Noradrenaline **F)** Adrenaline **G)** BI-167-107 association was read using nanoBRET between TS-SNAP-$\beta_2$AR-nLuc and Venus-mini-Gs which was read on PHERAstar FSX, at room temperature, using LUM 550LP/450-80nm module, $k_{obs}$ of $k_{slow}$ at each Venus-mini-Gs concentration was obtained by fitting association to a two phase association model, All figures show representative raw data of n=3

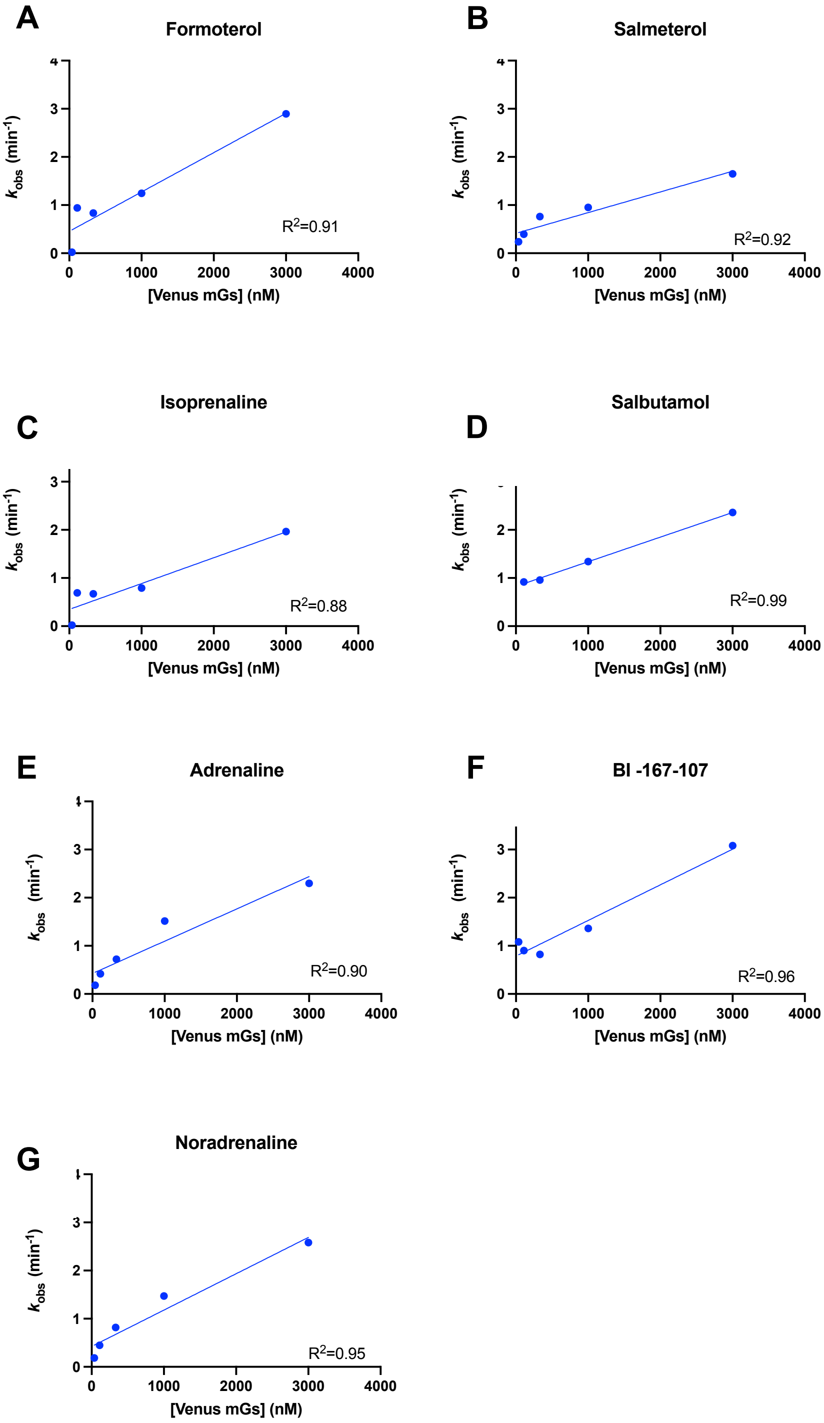


**SF6: *K*obs plots of *K*fast for Venus-mini-Gs association to DDM solubilised TS-SNAP-$\beta_2$AR-nLuc bound to A)** Formoterol **B)** Salmeterol **C)** Isoprenaline **D)** Salbutamol **E)** Adrenaline **F)** BI-167-107 **G)** Noradrenaline, association was read using nanoBRET between TS-SNAP-$\beta_2$AR-nLuc and Venus-mini-Gs which was read on PHERAstar FSX, at room temperature, using LUM 550LP/450-80nm module, *k*obs of *k*fast at each Venus-mini-Gs concentration was obtained by fitting association to a two phase association model, All figures show representative raw data of n=3, fitted to a linear model.

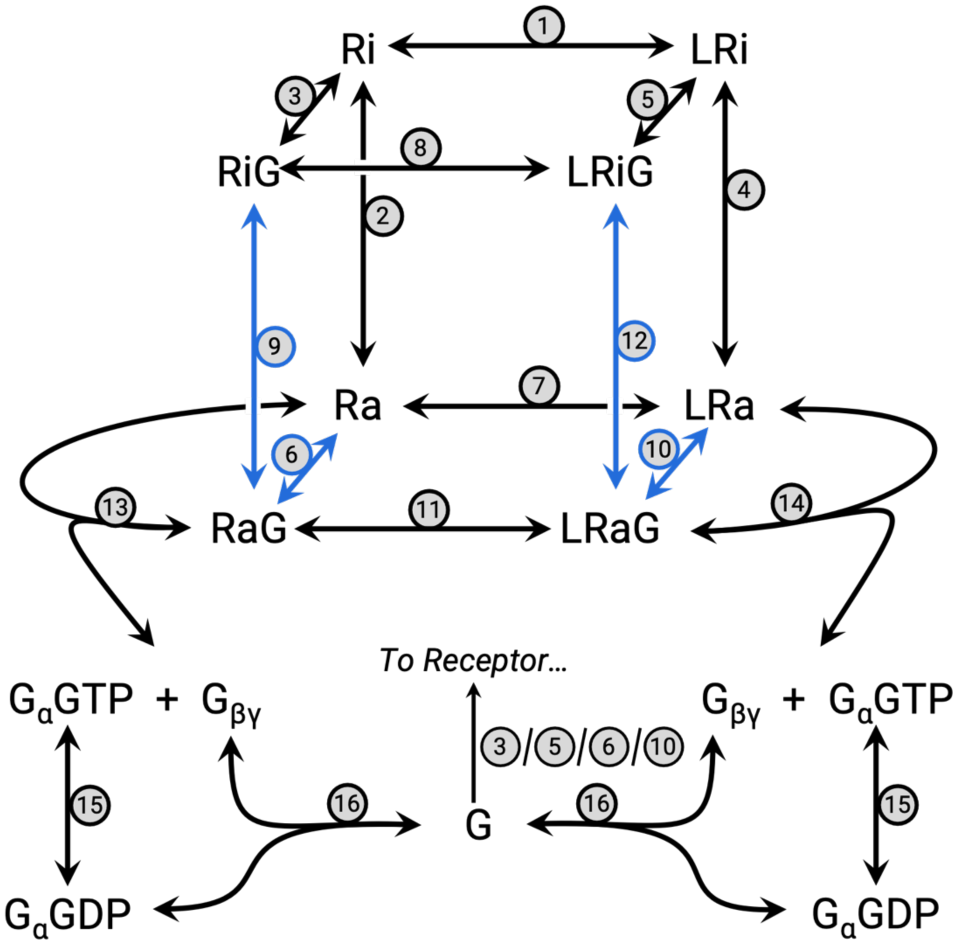


**SF7: Schematic of cubic ternary complex model.** A cubic ternary complex model with extended reactions enabling simulation of Gα activation cycle (Biomodels ID:2306220001). Base parameters are shown in **ST4**. To simulate changes in rates of G protein recruitment to active receptor the term $\beta^+$ was varied over a range of $10^{-2}$ to $10^4$. Reactions varied by changing the $\beta^+$ are shown in blue.

**ST1:** A summary of the saturating concentrations of $\beta_2$AR agonists used in in-solution TS-SNAP-$\beta_2$AR-Venus-mini-$G_s$ nanoBRET binding assays.

| Ligand | p$K_i$ | Saturating concentration used |
|---|---|---|
| **Formoterol** | 7.8 (15nM) | 5μM |
| **Isoprenaline** | 6.4 (398nM) | 100μM |
| **Adrenaline** | 5.2 (6309nM) | 500μM |
| **Noradrenaline** | 4.4 (39811nM) | 1mM |
| **BI-167-107** | 9.2 (0.63) | 100nM |
| **Salmeterol** | 9.1 (0.79) | 50nM |
| **Salbutamol** | 5.8 (1585nM) | 60μM |

**ST2: Quantification of the percentage of** Venus-mini-$G_s$ binding DDM-TS-SNAP-$\beta_2$AR that could be attributed to the fast association phase at varying [Venus-mini-$G_s$] and in the presence of each $\beta_2$AR agonist used in this study.

| | **% $k_{fast}$ at varying [Venus-mini-Gs] (nM)** | | | |
|---|---|---|---|---|
| | [3000] | [1000] | [333] | [111] |
| Formoterol | 71.4±3.0 | 63.6±6.5 | 49.9±11.5 | 23±9.0 |
| Isoprenaline | 60.0±12.1 | 53.5±10.8 | 41.6±17.7 | 23±11.0 |
| Salbutamol | 66.4±1.6 | 44.0±9.7 | 39.6±15.2 | 52±21.0 |
| C26 | 78.7±1.4 | 63.6±3.4 | 48.3±7.0 | 27±11.4 |
| Adrenaline | 76.5±3.9 | 66.3±8.1 | 45.6±11.9 | 35±13.8 |
| BI-167-107 | 62.9±13.0 | 66.1±3.6 | 51.4±3.2 | 43±8.5 |
| Salmeterol | 70.0±.3.8 | 48.6±9.1 | 34.6±9.8 | 36 ±15.3 |

Data are mean of n=3-4 experiments ±SEM.

**ST3:** A summary of the percentage of DDM-TS-SNAP-$\beta_2$AR: Venus-mini-$G_s$ complexes that dissociated with each $\beta_2$AR agonist at each concentration.

| | **% Dissociated** |
|---|---|
| **Formoterol** | 74.3±4.1 |
| **Isoprenaline** | 73.1±2.1 |
| **Salbutamol** | 72.3±2.2 |
| **C26** | 76.9±1.4 |
| **Adrenaline** | 73.1±2.1 |
| **BI-167-107** | 79.7±4.5 |
| **Salmeterol** | 72.5±1.5 |

Values are mean of n=3-4 experiments ±SEM.

**ST4: Base parameter sets for computational simulations.** Initial values and descriptions for each reaction parameter and starting species.

| Label | Description | Value | Units |
|---|---|---|---|
| $k_{L+}$ | Ligand binding rate | 1.00E+05 | $M^{-1}$ $s^{-1}$ |
| $k_{L-}$ | Ligand unbinding rate | 1.00E-02 | $s^{-1}$ |
| $k_{act+}$ | Receptor activation rate | 3.00E-02 | $s^{-1}$ |
| $k_{act-}$ | Receptor deactivation rate | 1.00E+02 | $s^{-1}$ |
| $k_{G+}$ | G protein binding rate | 1.00E+05 | $M^{-1}$ $s^{-1}$ |
| $k_{G-}$ | G protein unbinding rate | 1.10E-02 | $s^{-1}$ |
| $\alpha_+$ | Forward cooperativity factor for ligand bound receptor activation | 3.00E+03 | - |
| $\alpha_-$ | Backwards cooperativity factor for ligand bound receptor activation | 1.00E-01 | - |
| $\beta_+$ | Forward cooperativity factor for G protein-bound receptor activation | 1.00E+01 | - |
| $\beta_-$ | Backwards cooperativity factor for G protein-bound receptor activation | 1.00E+00 | - |
| $\gamma_+$ | Forward cooperativity factor for ligand binding a G protein-bound receptor | 1.00E+00 | - |
| $\gamma_-$ | Backwards cooperativity factor for ligand binding a G protein-bound receptor | 1.00E+00 | - |
| $\delta_+$ | Forward cooperativity factor for ligand bound, G protein-bound receptor activation | 1.00E+00 | - |
| $\delta_-$ | Backwards cooperativity factor for ligand bound, G protein-bound receptor activation | 1.00E+00 | - |
| $k_{GDA+}$ | G protein dissociation rate from active, G protein-bound receptor (possibly ligand-bound) | 1.00E+06 | $s^{-1}$ |
| $k_{GDA-}$ | Reformation of active, G protein-bound receptor (possibly ligand-bound) | 1.00E-10 | $M^{-2}$ $s^{-1}$ |
| $k_{GRA+}$ | Heterotrimeric G protein reassociation rate | 1.00E+08 | $M^{-1}$ $s^{-1}$ |
| $k_{GRA-}$ | G protein spontaneous dissociation rate | 1.00E-10 | $s^{-1}$ |
| $k_{hyd+}$ | Rate of hydrolysis of GαGTP | 1.00E-02 | $s^{-1}$ |
| $k_{hyd-}$ | Spontaneous exchange rate of GDP for GTP | 2.00E-06 | $s^{-1}$ |
| $[Ri]_{initial}$ | Initial concentration of inactive Receptor | 4.15E-10 | M |
| $[G]_{initial}$ | Initial concentration of heterotrimeric G protein | 4.15E-10 | M |
| $[L]_{tot}$ | Total concentration of Ligand | Varies | M |
